# Task-guided Generative Adversarial Networks for Synthesizing and Augmenting Structural Connectivity Matrices for Connectivity-Based Prediction

**DOI:** 10.1101/2024.02.13.580039

**Authors:** Tatsuya Yamamoto, Tomoki Sugiura, Tomoyuki Hiroyasu, Satoru Hiwa

**Affiliations:** Graduate School of Life and Medical Sciences, Doshisha University. (1-3 Tatara Miyakodani, Kyotanabe, Kyoto 610-0394, Japan); Department of Biomedical Sciences and Informatics, Doshisha University. (1-3 Tatara Miyakodani, Kyotanabe, Kyoto 610-0394, Japan)

**Keywords:** Data augmentation, structural connectivity, generative adversarial networks (GANs), connectome-based prediction, generative models

## Abstract

Recent machine learning techniques have improved connectome-based predictions by modelling complex dependencies between brain connectivity and cognitive traits. However, they typically require large datasets that are costly and time-consuming to collect. To address this, we propose Task-guided GAN II, a novel data augmentation method that uses generative adversarial networks (GANs) to expand sample sizes in connectome-based prediction tasks. Our method incorporates a task-guided branch within the Wasserstein GAN framework, specifically designed to synthesize structural connectivity matrices and improve prediction accuracy by capturing task-relevant features. We evaluated Task-guided GAN II on the prediction of fluid intelligence using the NIMH Health Research Volunteer Dataset. Results showed that data augmentation improved prediction accuracy. To further assess whether augmentation can substitute for increasing real-world sample sizes, we conducted additional validation using the HCP WU-Minn S1200 dataset. Task-guided GAN II improved prediction performance with limited real data, with gains of up to twofold augmentation observed. However, excessive augmentation did not result in further improvements, suggesting that augmentation complements, but does not fully replace, real data augmentation. These results suggest that Task-guided GAN II is a promising tool for harnessing small datasets in human connectomics research, improving predictive modelling where large-scale data collection is impractical.

## 1. Introduction

Connectomics, which represents neural circuits as networks and analyzes their topology through mathematical approaches, has emerged as an essential methodology in cognitive and computational neuroscience [1], [2]. It facilitates a deeper understanding of the complex relationships among brain networks, cognition, behavior, and individual differences in human cognitive traits and behaviors. Additionally, recent advances in machine learning methodologies have employed connectome-based machine learning models using structural and functional connectivity matrices to predict behavior or cognitive traits [3].

Recent studies have revealed that building prediction models requires more than a hundred samples [4], [5] and has also been shown to be improved in accuracy and robustness as the sample size becomes larger, not depending on the choice of regression models [6]. Therefore, researchers have often used large-sample open datasets such as the Human Connectome Project (HCP) Dataset (*N* = 1200) [7], ABCD Study (*N* = 11878) [8], UK Biobank (*N* = 40000+) [9]. These datasets have contributed significantly to developing analytical methods and understanding human connectomics [10]. However, Yeung et al. have reported that most self-recruited samples included around 100 subjects [5]. In the exploratory stage of the research, it is not easy for individual researchers to collect large samples over 100 due to the limitations on time and cost.

One of the best solutions to resolve the small sample size issue is to employ a data augmentation that generates a new sample by manipulating the given small dataset. In traditional data augmentation, geometric transformation has often been utilized in image data, such as rotation or flipping to the original data. However, such manipulations are inapplicable to the matrix-shaped data of brain functional and structural connectivity because their shape was constrained by the order of brain regions. In recent years, data augmentation methods using generative models have been proposed [11], [12], [13], [14], [15], [16], [17]. In these methods, generative augmentation models approximate the distribution of a given dataset and synthesize new samples with similar characteristics to the original dataset. Specifically, generative adversarial networks (GAN) have been increasingly used to data synthesis in numerous application fields with its strong feature of learning the local and global structure of data [18]. In the neuroscience field, Chao Li et al. proposed the structural connectivity augmentation method named BrainNetGAN to build the model for the classification of Alzheimer’s disease [15]. The accuracy of the graph neural network-based classification model of Alzheimer’s disease was improved using the structural connectivity data synthesized by their BrainNetGAN. Ruizhe Li et al. also suggested a data augmentation method for T1-weighted image named Task-guided GAN (TG GAN) for the brain age prediction model [16]. They improved the accuracy of the age prediction [16]. In their TG GAN model, the task-guided branch of a regression model for brain age prediction was incorporated into the GAN architecture. This accelerated the data synthesis to be more task-specific and contributed to improving the prediction task.

There are a lot of studies on data augmentation methods for medical images, but a few studies on brain connectome data augmentation methods for the connectome-based prediction task. The BrainNetGAN, as mentioned above, was developed for the classification task and trained using specific class labels to synthesize conditioned data. On the other hand, the TG GAN was designed with a 3D image input structure specifically for the T1-weighted image synthesis. Neither model is suitable for synthesizing connectivity matrices aimed at prediction tasks. Given the rapidly increasing interest in connectome-based prediction studies, there is an urgent need to develop a data augmentation model tailored to these tasks. Here, we propose a novel data augmentation method, Task-guided GAN II (TG GAN II). It was designed for synthesizing the structural connectivity matrices and employs a task-guided branch to predict human cognitive traits from the structural connectivity. We aim to enhance the accuracy of predicting cognitive traits from structural connectivity by augmenting the original dataset with the data synthesized by our TG GAN II. To assess the effectiveness of our method, we utilize the NIMH Health Research Volunteer Dataset [19] and the HCP WU-Minn S1200 release [7] to build the model predicting fluid and crystallised intelligence based on structural connectivity. The TG GAN II employs the Wasserstein GAN (WGAN) without task-guided branches as its baseline model. It is evaluated against this baseline in the following three perspectives: 1) the similarity of graph features between synthesized and original data, 2) the improvement in prediction performance, and 3) the influence of the added synthetic data volume on prediction performance enhancement. Furthermore, the effect of data augmentation using TG GAN II on prediction accuracy was evaluated by comparing it with the effect of increasing the sample size using real data. We hope that these investigations will make a substantial contribution toward addressing the sample size challenges in connectome-based prediction tasks.

## 2. Methods

### 2.1 Baseline model: Wasserstein GAN with gradient penalty (WGAN-GP)

We employed a Wasserstein GAN with gradient penalty (WGAN-GP) [20] as the baseline model for our proposed TG GAN II. The Wasserstein GAN incorporates the Wasserstein distance (also known as the Earth Mover’s Distance) into its objective function instead of the Kullback-Leibler (KL) divergence, which is commonly used in traditional GANs. This modification helps improve model stability and mitigates mode collapse. Both the Wasserstein distance and KL divergence are measures of similarity between two probability distributions. In the context of GANs, these distributions correspond to the real and generated data. The Wasserstein distance can be understood as the minimum effort required to transform one probability distribution into another. This concept is analogous to redistributing a pile of dirt to match the shape of another pile, where the cost is determined by the amount of dirt moved and the distance it is transported. This property enables more stable gradient updates, thereby addressing common issues in GAN training. The WGAN-GP consisted of the generator that synthesized new structural connectivity matrices, and the critics assessed the synthesized images by estimating the Wasserstein distance between synthesized and given ones. The gradient penalty is used to enforce a Lipschitz constraint on the critics. We used an autoencoder-based generator in our WGAN-GP model to synthesize new images by interpolating the latent feature space (**Figure 1**).

**Figure 1.**
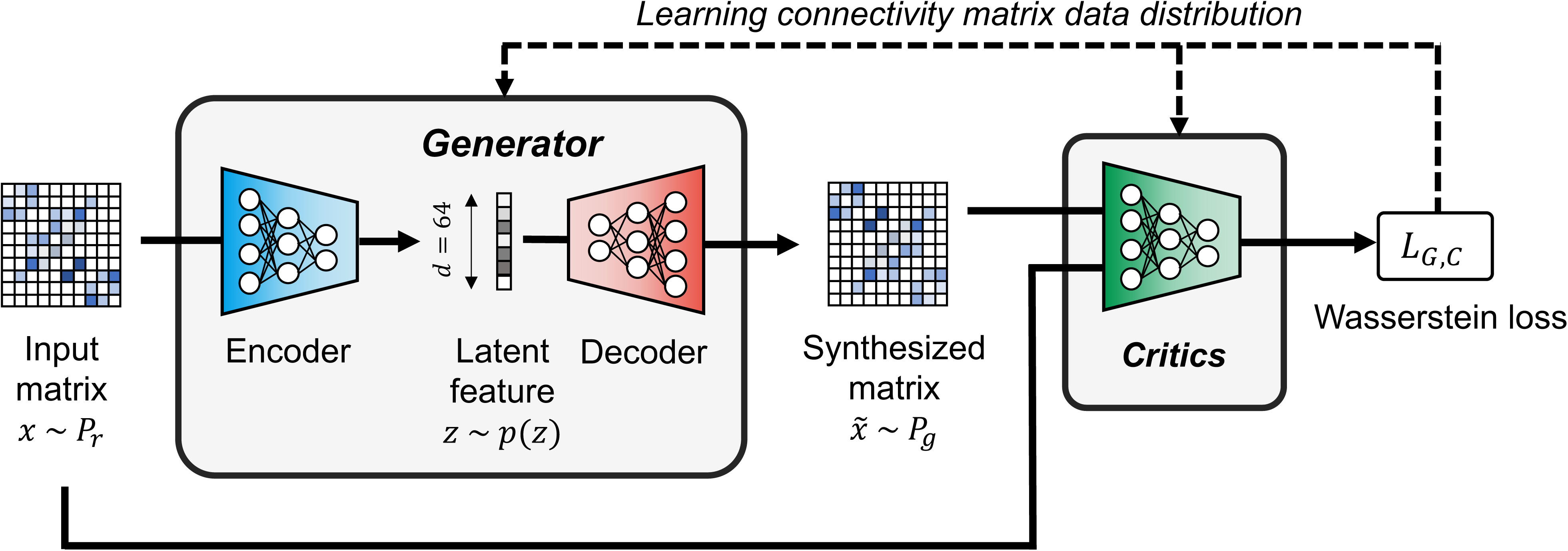
The architecture of the WGAN-GP used for synthesizing connectivity matrices, serving as the baseline model, where *P*_*r*_ and *P*_*g*_ denote the distributions of given and synthesized connectivity matrices, respectively, and *x* and 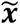 are the given input and synthesized matrices, respectively. The encoder transforms the input matrix into latent variables *z*, while the decoder reconstructs the matrix 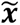 from *z*. The critics aim to estimate the Wasserstein distance between the given and synthesized matrices, while the generator seeks to minimize this distance.

The encoder in the generator module extracted features from given connectivity matrices and mapped them into the latent space, and the decoder module synthesized the matrices from the latent variables. We employed convolutional neural networks for brain networks (BrainNetCNN) proposed by Kawahara et al. [21], which consisted of four convolutional layers and two fully connected layers for the encoder. The BrainNetCNN had three convolutional layers specialized for extracting topological features of connectivity matrices: edge-to-edge (E2E), edge-to-node (E2N), and node-to-graph (N2G) convolutions (**Figure 2**). We adopted a simple feed-forward neural network with five fully connected layers for the decoder. Besides, we used the softplus function as an activation function in the final layer because the elemental value of structural connectivity took zero or more. The decoder outputted a vector of the lower triangle part in the connectivity matrix, which was finally matricized. We also utilized the BrainNetCNN architecture with four convolutional layers for the critics.

**Figure 2.**
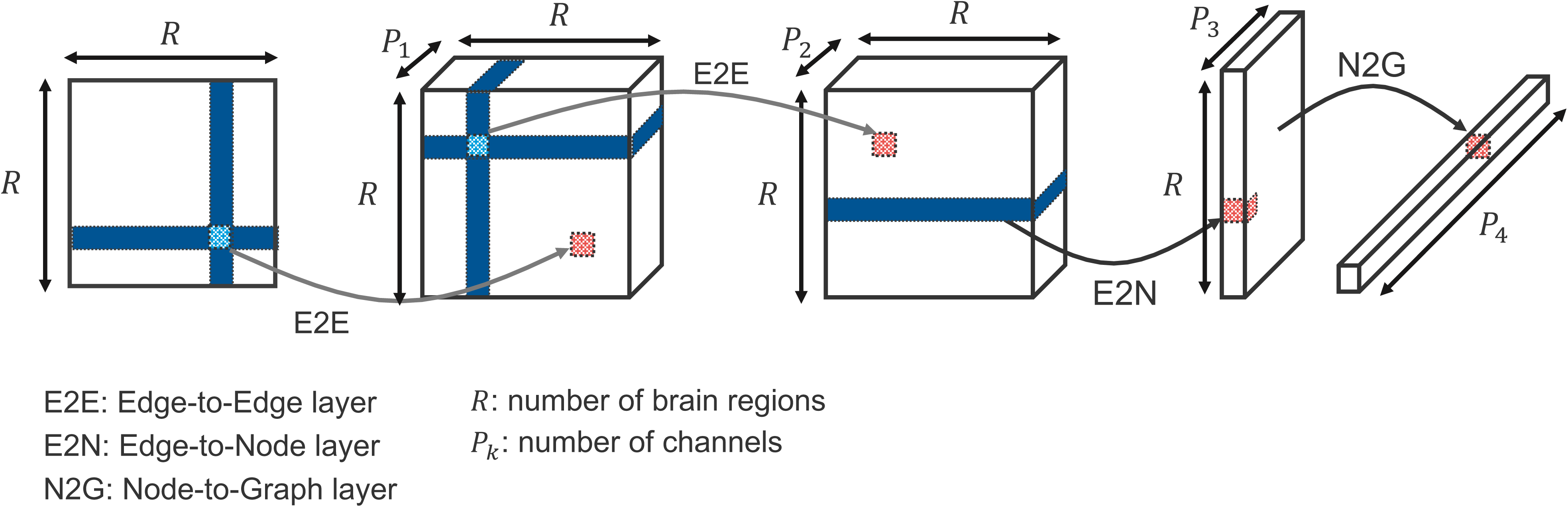
Architecture of BrainNetCNN. This architecture introduces convolution kernels designed for the adjacency matrix of brain networks, a development unique to BrainNetCNN. These kernels are applied to an element’s entire row and column, enabling edge-to-edge (E2E) and edge-to-node (E2N) convolution operations. Initially, the connectivity matrix undergoes convolution with one or more E2E kernels, emphasizing the connections between adjacent brain regions. Subsequently, the output from the E2E convolution is processed through an E2N filter, which calculates a weighted sum of edges for each brain region. A node-to-graph (N2G) kernel integrates these weighted node values to produce a single output value, representing an aggregate measure of brain connectivity.

The mathematical description of the loss function for the generator and critics in our WGAN-GP is presented in Equation 1. The role of the critics is to evaluate the similarity between the synthesized and acquired matrices by estimating the Wasserstein distance. To achieve this, the critics maximize the objective function, effectively increasing the Wasserstein distance to provide a more precise measure of dissimilarity. This process guides the generator in minimizing the distance, thereby making the synthesized matrices progressively more similar to the acquired ones. Conversely, the generator was trained to minimize the same equation, bringing the distributions of the synthesized and original matrices closer together.

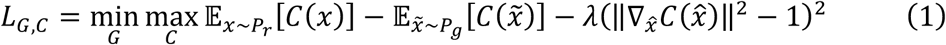

where *P_r_* is the distribution of given connectivity matrices, and *P_g_* is the distribution of synthesized matrices. *C*(*x*) denotes the output of the critics for a given input *x*, and 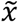 represents the output of the decoder *G*(*z*) when given a latent variable *z*∼*p*(*z*) generated in the encoder. The third term in the Equation 1 introduces a gradient penalty for encouraging *C*(*x*) to adhere to the 1-Lipschitz continuity, with *λ* being the coefficient of this penalty.

### 2.2 Task-guided GAN II

In this study, we have built upon the baseline WGAN-GP model to develop a novel GAN-based connectivity synthesis model named TG GAN II. Our proposed model extends the WGAN-GP by incorporating a task-guided branch specifically designed for prediction tasks. Its objective is to augment the dataset with additional connectivity matrices and their corresponding objective variables, thereby facilitating the construction of a more accurate cognitive trait prediction model based on brain connectivity.

**Figure 3** illustrates the architecture of TG GAN II. Within this framework, a regressor model for cognitive trait prediction was integrated into the WGAN-GP structure as the task-guided branch to enhance the performance of the prediction task. This branch was designed to generate a latent space that captured the variations in both explanatory and objective variables. It consisted of four convolutional layers and four fully connected layers, culminating in the output of the predicted objective variable from the synthesized connectivity matrix. The loss function for the task-guided regressor of TG GAN II, outlined in Equation 2, was calculated by the root mean squared error (RMSE) between the observed and predicted objective variables, with both the generator and critics optimized to minimize this function:

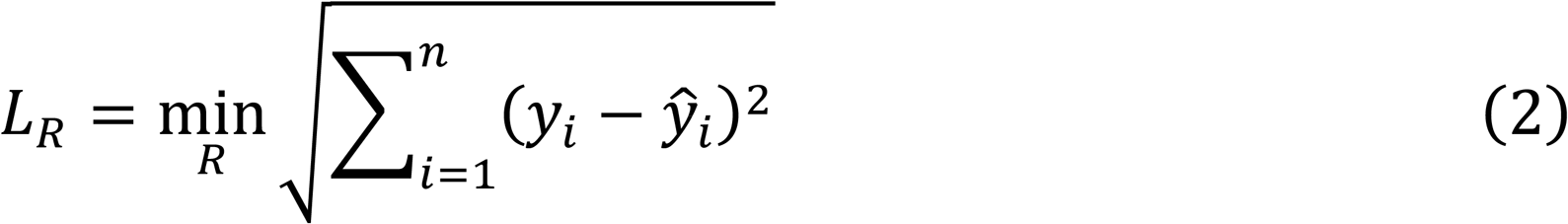

where *y_i_* denotes the observed objective variable and 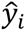 is the predicted one. In the training of TG GAN II, both Equations 1 and 2 were optimized simultaneously, resulting in the comprehensive loss function for TG GAN II being represented in Equation 3:

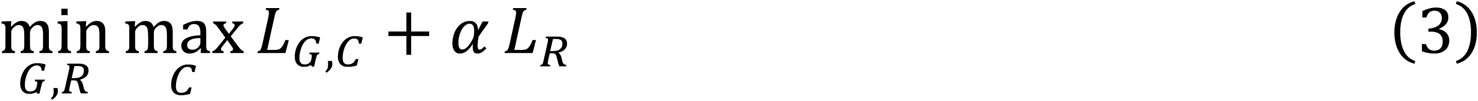

where the coefficient *α* served as the weight against *L_R_*, the regression loss, balancing the two terms of the loss function.

**Figure 3.**
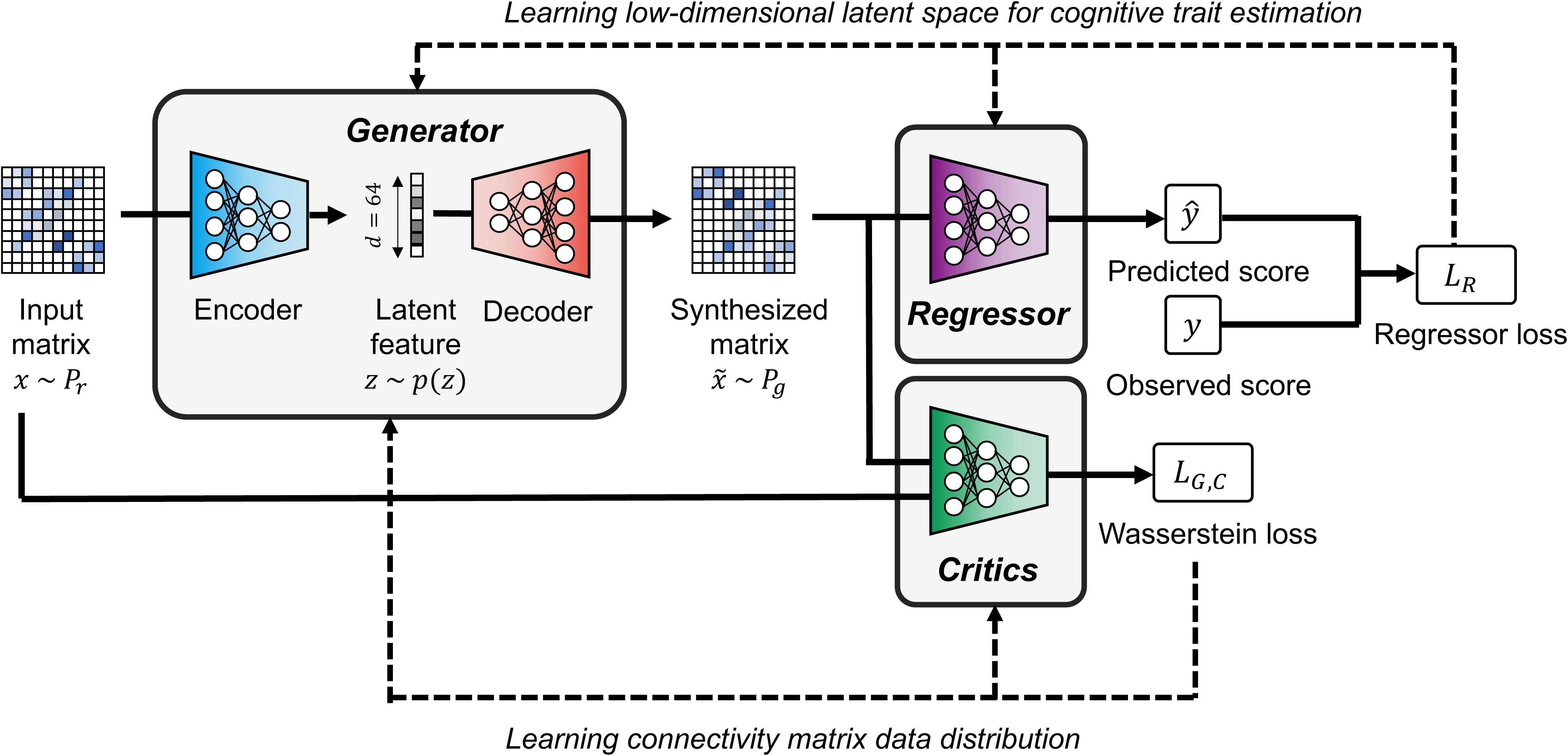
The architecture of Task-guided GAN II for synthesizing connectivity matrices. This model introduces modifications over the WGAN-GP, notably incorporating a task-guided regressor branch and specialized loss function calculations. The regressor branch features four convolutional layers and four fully connected layers, which together output the predicted objective variable based on the synthesized connectivity matrix. The loss function of the WGAN-GP is expanded to include the root mean squared error (RMSE) between the observed and predicted objective variables, facilitating more accurate predictions.

### 2.3 Latent space interpolation for connectivity matrices and objective variable synthesis

Using the trained TG GAN II model, we aimed to synthesize new connectivity matrices and their corresponding objective variables by performing the interpolation in the latent space. A single pair of the connectivity matrix and its objective variable was synthesized using two pairs of them (*X_i_*, *y_i_*) and (*X_j_*, *y_j_*) following Equations 4 and 5 referring to the method provided by Li et at. [16]:

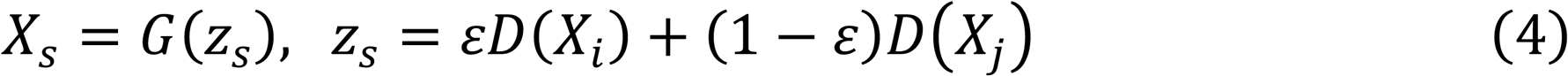

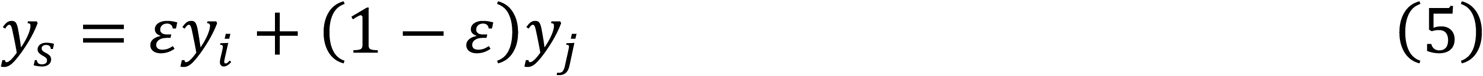

where *ε* is the value between 0 and 1. The value of *ε* was assigned in equal increments based on the total number of synthesized samples, ensuring a uniform interpolation between the selected pairs. This procedure is illustrated in **Figure 4**.

**Figure 4.**
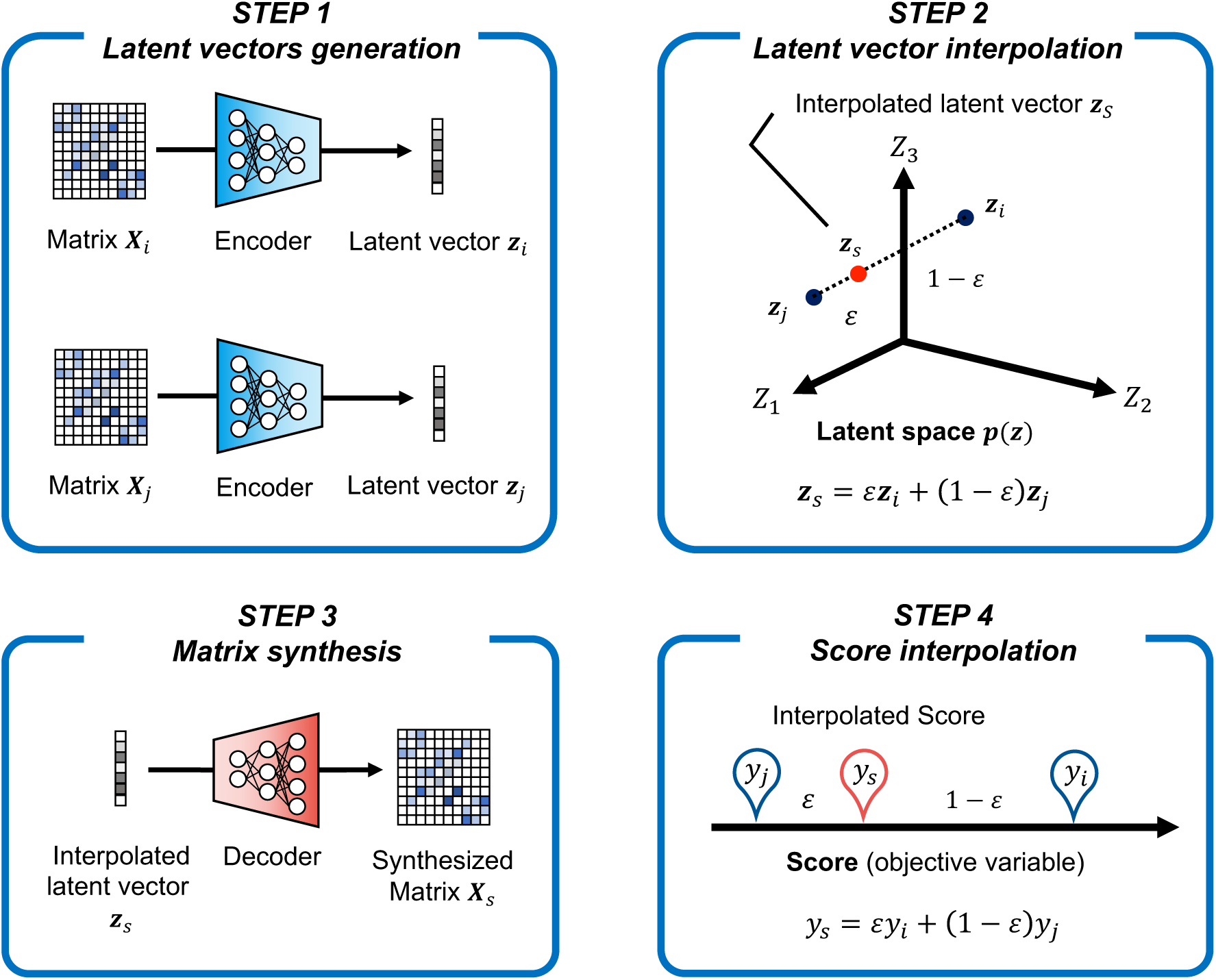
Synthesis of connectivity matrices and their corresponding objective variables through latent vector interpolation. This process involves using two pairs of given connectivity matrices and objective variables, (*X*_*i*_, *y*_*i*_) and (*X*_*j*_, *y*_*j*_), as inputs. These samples are first mapped to the latent space using the trained generative model. A new set of latent variables is then generated by performing linear interpolation between these mapped points. The interpolated latent variables are fed into the decoder, which reconstructs the connectivity matrices.

In Li et al., sample pairs (*i*, *j*) were selected under the constraint that their age difference was less than 2 years, ensuring that latent space variations were learned within similar age groups. Since the objective variables in our study differ from those in Li et al., we followed a similar approach but adapted it to our dataset characteristics. Specifically, we defined neighboring samples based on cognitive scores, treating adjacent (i.e., consecutive) samples in the dataset as neighbors. This approach assumes that locally, cognitive traits vary in an approximately linear manner, allowing for meaningful interpolations in the latent space.

### 2.4 Structural Connectivity Mapping

#### 2.4.1 Dataset

##### Dataset 1

In this study, we used MRI images and cognitive scores in the NIMH Healthy Research Volunteer Dataset (https://openneuro.org/datasets/ds004215/versions/1.0.1) [19] to evaluate the effectiveness of data augmentation using TG GAN II on prediction model performance. We used T1-weighted images, diffusion-weighted images, and the score of NIH Toolbox Cognition Battery [22] (age-adjusted score) as cognitive traits from 108 samples in which T1-weighted image, DWI, and resting fMRI had been obtained without deficiencies. We showed the average and standard variation of these scores in Table 1. We summed up four task scores in each participant and used it as a fluid intelligence score. This composed score shows a higher signal-noise ratio than each task score and is less affected by the variability of each task score [23]. In this study, Dataset 1 was used for in-depth analyses, including latent space exploration, network property comparisons between synthesized and acquired data, and the overall impact of data augmentation on prediction models.

**Table 1.** NIH Toolbox Cognition Battery Scores in the NIMH Healthy Research Volunteer Dataset.

| Domain | Average | SD |
| --- | --- | --- |
| Flanker Inhibitory Control and Attention | 91.3 | 12.1 |
| List Sorting Working Memory | 103.5 | 13.0 |
| Picture Sequence Memory | 114.9 | 16.7 |
| Dimensional Change Card Sort | 109.6 | 17.9 |
| Fluid Intelligence | 419.2 | 40.9 |

##### Dataset 2

Additionally, the HCP WU-Minn S1200 release was utilized to evaluate whether the synthesized data is substitute for actual data. Similarly to the NIMH Health Research Volunteer Dataset, we filtered participants based on the availability of a complete set of T1-weighted image, DWI, and the cognitive traits in the NIH Toolbox

Cognition Battery. Finally, we used 963 participants, for whom the structural connectivity matrix had been reconstructed. The average and standard variation of cognitive traits are shown in **Table 2**. From these scores, we summed up five task scores (Dimensional Change Card Sort, Flanker Inhibitory Control and Attention, Picture Sequence Memory, List Sorting Working Memory, and Pattern Comparison Processing Speed) and two task scores (Picture Vocabulary, and Oral Reading Recognition) in each participant and obtained the fluid and crystallised intelligence scores. Unlike Dataset 1, which was used for latent space analysis and detailed network property analyses, Dataset 2 was specifically employed to compare the effects of data augmentation against the effects of increasing the actual sample size. The larger sample size of Dataset 2 against Dataset 1 allowed us to examine whether a smaller dataset enhanced with synthetic data could achieve prediction accuracy comparable to a larger real dataset (e.g., N=100 with augmentation vs. N=200, 300, 400 real samples).

**Table 2.** NIH Toolbox Cognition Battery Scores in the HCP WU-Minn S1200 release.

| Domain | Average | SD |
| --- | --- | --- |
| Dimensional Change Card Sort | 101.9 | 10.1 |
| Flanker Inhibitory Control and Attention | 101.4 | 10.1 |
| Picture Sequence Memory | 104.7 | 16.8 |
| List Sorting Working Memory | 103.1 | 13.3 |
| Pattern Comparison Processing Speed | 103.0 | 19.9 |
| Fluid Intelligence | 514.0 | 45.7 |
| Picture Vocabulary | 108.6 | 15.7 |
| Oral Reading Recognition | 106.3 | 15.2 |
| Crystallised Intelligence | 214.8 | 28.5 |

#### 2.4.2 MRI data preprocessing

For the NIMH Health Research Volunteer Dataset, we preprocessed DWI and T1-weighted images using a pre-configured DWI process pipeline, QSIPrep (0.15.4) [24]. The preprocessing process consists of the following six steps: (1) T1-weighted image preprocessing, (2) Denoising DWI using Marchenko-Pastur Principal Component Analysis (MP-PCA) [25], (3) B1 field inhomogeneity correction, (4) Susceptibility distortion correction (TOPUP) [26], (5) Eddy current and head motion correction (EDDY) [27], (6) Registration of DWI into T1-weighted image.

We also preprocessed DWI and T1-weighted images using the MRtrix3_connectome pipeline (https://github.com/BIDS-Apps/MRtrix3_connectome), which is a toolbox configured based on the MRtrix3 software packages for the HCP WU-Minn S1200 release. The preprocessing process includes the six steps: (1) T1-weighted image preprocessing, (2) Denoising DWI using Marchenko-Pastur Principal Component Analysis (MP-PCA) [25], (3) Gibbs ringing artefacts removal, (4) B1 field inhomogeneity correction, (5) Susceptibility distortion correction (TOPUP) [26], (6) Eddy current and head motion correction (EDDY) [27], (7) Registration of DWI into T1-weighted image.

#### 2.4.3 Structural connectivity mapping

We also reconstructed whole brain tractograms and structural connectivity using QSIPrep (0.15.4) [24] for the NIMH Health Research Volunteer Dataset. The reconstruction of structural connectivity maps was performed in the following steps. First, the fiber orientation distribution function (fODF) was estimated using the Single-Shell 3-Tissue constrained spherical deconvolution (SS3T CSD) model [28] (the response function was estimated following Dhollander et al. [29]). Second, the whole brain tractogram was estimated to be 10 million streamlines using probabilistic (iFOD2 [30]) and anatomically-constrained tractography (ACT) [31]. Streamlines were seeded from a boundary between the white and gray matter derived from the structural segmentation. Third, we computed streamline weights based on the SIFT2 algorithm that reduced the biases in probabilistic tractography [32]. Finally, the whole brain was parcellated into 116 regions based on automated anatomical labeling (AAL), and the structural connectivity was defined as the sum of SIFT2-weighted streamlines connecting two arbitrary regions divided by the sum of the volumes of those regions.

For the HCP WU-Minn S1200 release, we reconstructed structural connectivity matrices similarly above utilizing MRtrix3_connectome. The structural connectivity matrices were reconstructed in the following steps. First, the fODF was estimated using the Multi-Shell Multi-Tissue CSD (MSMT CSD) model. Second, the whole brain tractogram was reconstructed to be 10 million streamlines using iFOD2 and ACT. Third, streamlines were weighted based on the SIFT2 algorithm. Finally, the whole cortex was delineated into 360 areas utilizing HCP MMP 1.0 atlas, and the structural connectivity was defined as described above.

### 2.5 Evaluation of the effect of Regressor branch on Task-guided GAN II (Dataset 1)

#### 2.5.1 Training setup for generative models

The entire dataset was divided into the discovery dataset (70%; *N* = 75) and the test dataset (30%; *N* = 33). In this division, we sorted the samples by their objective variables and then assigned the samples with a rank of (2nd, 3rd, 5th, 6th, …, 103rd, 105th, 106th, 108th) to the discovery dataset and (1st, 4th, 7th, …, 101st, 104th, 107th) to the test dataset. This data division method was proposed by Cui et al. [6], [33], and ensures a similar distribution of behavioral scores across datasets while minimizing random bias in the division. We adopted this method due to the small sample size of our dataset. The discovery dataset was used to train TG GAN II and WGAN-GP, select the data augmentation model, and construct the prediction model. Conversely, the test dataset validated the prediction model’s accuracy. We adopted a hold-out validation scheme for the training and model selection processes in TG GAN II and WGAN GP. Similar to the discovery-test splitting, we sorted the discovery samples according to their objective variables. Then we assigned samples with a rank of (2nd, 3rd, 5th, …, 72nd, 74th, 75th) to the training dataset and (1st, 4th, …, 73rd) to the validation dataset.

The TG GAN II and WGAN-GP models were trained with Adam optimizer with momentum parameters *β*_2_ = 0.9, *β*_2_ = 0.999 (learning rate: 0.0001, batch size: 2), and were run for 2000 epochs. We implemented an early stopping strategy to prevent overfitting on the training dataset. The training was stopped if the regressor loss on the validation dataset did not improve for 50 consecutive epochs after reaching the 500th epoch.

There are several hyperparameters in TG GAN II and WGAN-GP, including the dropout rate *p* and the coefficient *α*. The dropout rate *p* for the encoder, critics, and regressor was selected from the following set of values: 0.1, 0.2, 0.3, 0.4, and 0.5. Similarly, the coefficient *α* was chosen from a range of 0.1 to 1.0 in increments of 0.1. To optimize these hyperparameters, we performed a grid search, selecting the values that minimized the sum of the regressor loss on the training and validation datasets within the hold-out validation framework. For WGAN-GP models, which do not include a regressor, the dropout rate was set to match that of the selected TG GAN II model.

**Table 3** presents the computational resources and training time required for TG GAN II and WGAN-GP. To measure training time, we recorded the duration of 100 epochs, as the total number of training epochs varied due to the early stopping strategy used in TG GAN II.

**Table 3.** Computational resources and the training time on TG GAN II and WGAN-GP.

|  | Task-guided GAN II | WGAN-GP |
| --- | --- | --- |
| CPU | Intel Xeon W-2265 3.50 GHz, 12 cores |  |
| GPU | NVIDIA GeForce RTX 3060 12 GB<br>NVIDIA GeForce RTX 4070 12 GB |  |
| Memory | 64 GB |  |
| Training time [s/100 epoch] | 361.991 | 336.738 |

The code used for training and evaluating the models in this study is available on GitHub at https://github.com/MIS-Lab-Doshisha/tg-gan2.

#### 2.5.2 Synthesized connectivity matrices evaluation based on graph theory metrics

We evaluated the similarity between the structural connectivity matrices synthesized by TG GAN II and WGAN-GP and the original matrices within the discovery dataset. First, we synthesized a connectivity matrix for each sample using these models. To quantify the differences between the acquired and synthesized matrices, we subtracted the synthesized matrices from their corresponding acquired matrices for each generative model. Then, we calculated the average difference matrix for each model. These averaged difference matrices were visualized and compared.

In the subsequent analysis, we quantitatively assessed the similarity between the acquired and synthesized matrices using graph theory metrics: connectivity strength, betweenness centrality, and clustering coefficient. For this purpose, we computed the average matrices for both acquired and synthesized connectivity matrices and binarized them according to edge density thresholds, ranging from 5% to 25% in increments of 5%. We then estimated the distribution of these three graph theory metrics for the acquired and synthesized averaged matrices. The similarity between these distributions was quantified using the Kullback-Leibler (KL) divergence.

Additionally, to see that each individually generated SC network has similar network features to real SC networks, we also computed network measures for each individual SC matrix rather than using averaged matrices. Specifically, we adopted five graph-theoretical measures: betweenness centrality, clustering coefficient, modularity, global efficiency, and local efficiency. For nodal measures, we first computed them for each brain region and then averaged across regions. To compare acquired and synthesized matrices, we conducted a paired analysis and assessed the distribution differences.

#### 2.5.3 Latent space evaluation

We assessed the relevance between the fluid intelligence scores and the latent space generated by TG GAN II and WGAN-GP encoders. Latent variables *z* were sampled separately from the training and validation dataset. We projected the acquired latent variables onto a two-dimensional space using principal component analysis (PCA) to visualize the latent space. Subsequently, we calculated the Pearson correlation between the first principal component (PC1) scores and the corresponding fluid intelligence scores.

#### 2.5.4 Performance evaluation on data augmentation for connectome-based prediction

To evaluate the effectiveness of TG GAN II, we built the regression model to predict the fluid intelligence scores from the structural connectivity matrices augmented by two generative models. We used the Ridge regression algorithm [34], which is widely used in the neuroscience field. The prediction performance of the fluid intelligence was then compared between models augmented by TG GAN II and WGAN-GP.

To assess the prediction model’s efficacy within the discovery dataset, we utilized repeated 5-fold cross-validation (5F CV). Additionally, the L2 regularization parameter in Ridge regression was fine-tuned using 5-fold cross-validation within each fold of the outer 5F CV. The methodology for this repeated nested CV process is outlined in subsequent sections.

##### Repeated outer 5F CV

In the outer 5F CV phase, the discovery dataset was randomly divided into five subsets. Four subsets were merged and used as the training dataset, while the remaining subset served as the validation dataset. We constructed the prediction model using the training dataset and the parameters determined during the inner 5F CV phase. This model was then validated against the validation dataset. This cycle of training and validation was repeated until each subset had been used as the validation dataset once. The prediction accuracy was calculated as the Pearson correlation between the observed and predicted scores. The outer 5F CV was repeated 20 times to avoid the bias resulting from random splitting.

##### Inner 5F CV and parameter tuning

Within each iteration of the outer 5F CV, the L2 regularization parameter for Ridge regression was optimized through an inner 5F CV process. We selected this parameter from a set of 16 possible values: *α* ∈ {*x* | *x* = 2*^n^*, *n* ∈ ℤ, *n* ∈ [−10, 5]}. For the inner 5F CV, the training dataset from the outer loop was randomly split into five subsets. Four subsets were used in each iteration to train the model under each parameter setting, with the fifth subset reserved for validation. This procedure was executed five times, ensuring each subset was utilized as the validation dataset. For each parameter across the inner 5F CV loops, we calculated a Pearson correlation and a mean absolute error (MAE) between the observed and predicted scores, averaging these metrics across all loops. The sum of the mean correlation and the inverse of MAE, standardized for differing scales, served as the measure of prediction accuracy in inner validation [35], [36]. The parameter that yielded the highest accuracy model was selected in the subsequent outer 5F CV.

##### Evaluation of out-of-sample prediction accuracy

To evaluate the model trained using repeated nested CV, 100 prediction models from the outer 5F CV were validated using the test dataset. Prediction accuracy was assessed using Pearson’s correlation and RMSE between observed and predicted scores.

##### Evaluation of prediction performance

We evaluated the accuracy of the fluid intelligence score prediction model augmented by TG GAN II and WGAN-GP. Data augmentation was applied to the discovery dataset, increasing the structural connectivity matrices and fluid intelligence scores to double (original data + 100%), triple (+ 200%), quadruple (+ 300%), and quintuple (+ 400%) through the latent space interpolation, as detailed in section 2.4.6. Utilizing the original and augmented datasets, we constructed fluid intelligence prediction models using the repeated nested CV method outlined in section 2.5.1. These models were then validated against the test dataset. To compare the prediction accuracy between the data-augmented models and the baseline (non-augmented: original sample size) model, we conducted Welch’s *t*-test. For this analysis, outlier detection was performed on the accuracy of 100 prediction models, identifying any values more than three standard deviations from the mean as outliers. Additionally, we applied the Bonferroni correction for multiple comparisons.

#### 2.5.5 Evaluation of reliability of estimated feature weights

In connectome-based prediction of cognitive traits, a key concern is the reliability of feature weights. To assess this, we calculated the mean and the 95% confidence interval (CI) range for each weight, based on coefficients obtained from 100 prediction models trained in the previous section. These two metrics were compared between prediction models trained with the augmented data using TG GAN II and WGAN-GP to evaluate the stability of feature importance across different methods.

### 2.6 Comparison of data augmentation and actual sample size increase (Dataset 2)

To evaluate whether data augmentation can serve as a substitute for increasing the actual dataset size, we conducted additional experiments using Dataset 2. Specifically, we compared the predictive performance of models trained with augmented data against those trained with progressively larger subsets of the HCP WU-Minn S1200 release dataset.

The entire dataset was divided into a subset for the generative model (N=321) and a subset for the prediction model (N=642). For generative model training, the dataset (N=321) was further split into a training set (N=241) and a validation set (N=80). For the prediction model, the dataset (N=642) was partitioned into a training set (N=482) and a test set (N=160).

The baseline sample size for the prediction model was set to N=80 out of the total 482 available samples in the training set, and synthesized samples generated by the trained generative model were added to the baseline with different augmentation sizes: N=161 (baseline +100%), N=242 (baseline +200%), N=322 (baseline +300%), N=402 (baseline +400%), and N=482 (baseline +500%).

For comparison, we trained the prediction model using the actual training dataset with the same sample size settings as the augmented dataset (N=80, 161, 242, 322, 402, 482). We constructed regression models using both fluid intelligence and crystallised intelligence as objective variables and evaluated their predictive performance. The effects of data augmentation were compared against those of increasing the actual dataset size in terms of prediction model performance, quantified by the RMSE of the prediction model and the Pearson correlation between actual and predicted objective variables.

For all generative and prediction model training, the assignment of objective variables to each sample followed the method used in Dataset 1. However, in Dataset 2, different hyperparameters and a regression algorithm were applied in generative and prediction model training to achieve sufficient prediction accuracy. For generative model trainings, the dropout rate *p* for the encoder, critics, and regressor was selected from the set of values: 0.1, 0.2, 0.3, 0.4, and 0.5. Similarly, the coefficient *α* was also chosen from the set of values: 0.1, 0.2, 0.3, 0.4, and 0.5. For prediction model training, we employed the Elastic-Net regression algorithm, which, like Ridge regression, is widely used in neuroscience. Elastic-Net regression algorithm is effective in high-dimensional datasets as it performs variable selection due to the L1-penalty. In this model, the regularization parameter *α* was chosen from the set of values: *α* ∈ {*x* | *x* = 2^2*n*^, *n* ∈ ℤ, *n* ∈ [−5, 4]}. Additionally, the relative weighting of L1 and L2-norm balance, *λ*, was selected from the values: 0.2, 0.4, 0.6, 0.8, and 1.0. Other training configurations were conducted following the procedures established in Dataset 1 to ensure consistency across experiments.

## 3. Results

### 3.1 Analysis of synthesized connectivity matrices (Dataset 1)

**Figure 5** presents the average connectivity matrices for the acquired and the synthesized structural matrices using TG GAN II and WGAN-GP, with brain regions aligned according to Yeo’s 7 Network definition. In **Figure 5A**, the weighted matrices are displayed, whereas **Figure 5B** showcases those binarized at a 25% edge density threshold. The average matrices in **Figure 5** were computed separately for the acquired and synthesized structural matrices. The synthesized matrices were generated by the trained Generator using the discovery dataset as input, and their average was taken over all synthesized samples. Similarly, the acquired matrices were averaged over the corresponding input matrices from the discovery dataset. This approach ensures consistency in comparison between the synthesized and acquired data. The synthesized matrices from WGAN-GP exhibit connections that do not exist in the acquired matrices and are not observed with TG GAN II. This is evident in both the weighted and binarized connectivity matrices.

**Figure 5.**
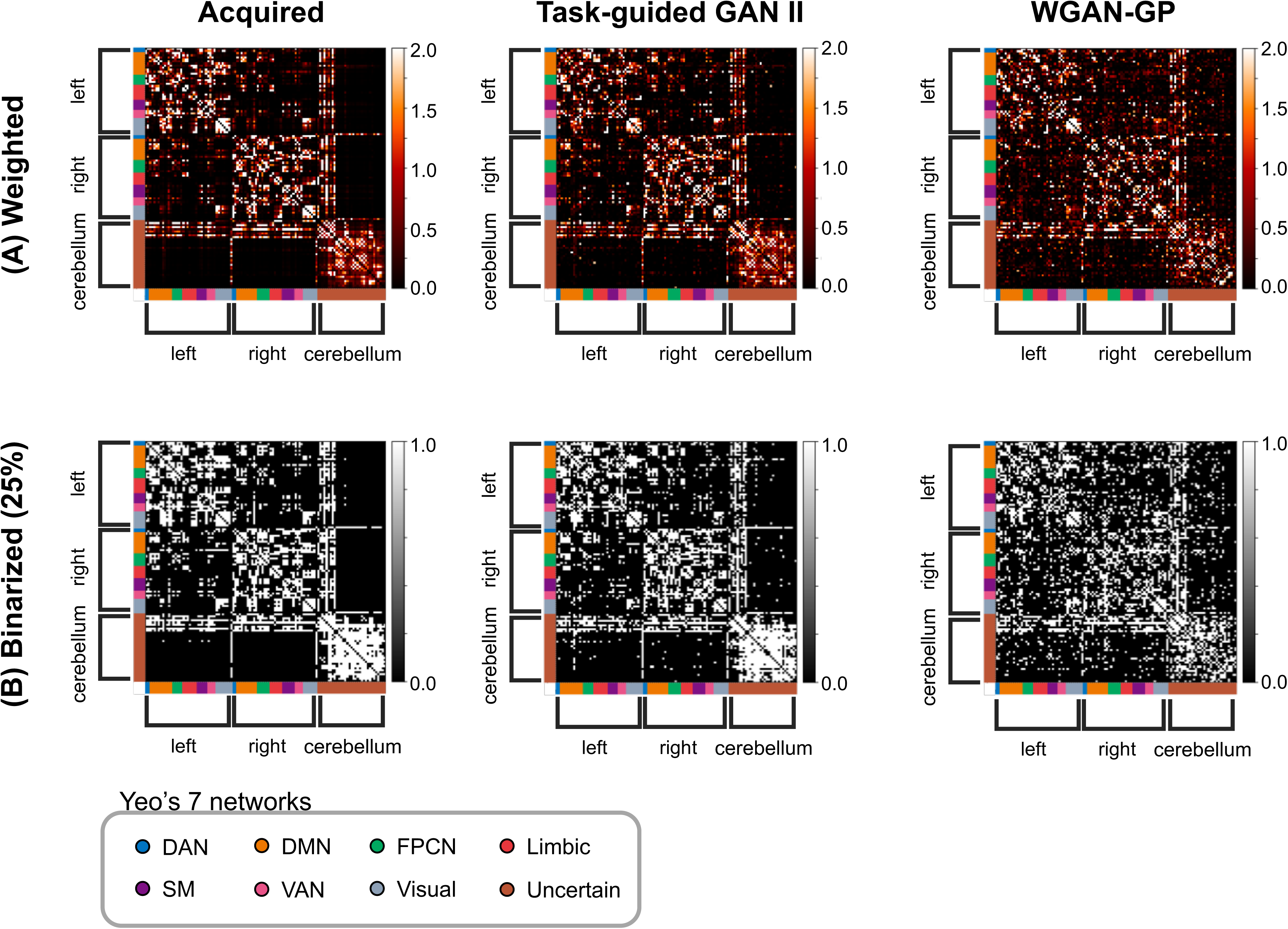
Comparison between acquired and synthesized structural connectivity matrices using Task-guided GAN II and WGAN-GP. (A) displays weighted connectivity matrices, and (B) shows binarized connectivity matrices, where binarization was applied to retain only the top 25% of the strongest connections. The brain regions in matrices were aligned according to Yeo’s 7 Network definition.

**Figure 6** displays the average difference in connectivity matrices between the acquired and those synthesized by TG GAN II and WGAN-GP. Each part of the connectivity matrix differentiated the connectomes: intra-hemispherical connections (left hemisphere: A, right hemisphere: C), inter-hemispherical connections (B), cerebrum-cerebellum connections (D), and intracerebellar connections (E). The differences were notable in the intrahemispheric and intracerebellar connections. TG GAN II and WGAN-GP tended to estimate weaker connections than those observed in the actual data. Additionally, WGAN-GP was shown to generate more interhemispheric and cerebrum-cerebellum connections compared to TG GAN II, indicating that TG GAN II synthesized connectivity matrices similar to the acquired data in these regions than WGAN-GP.

**Figure 6.**
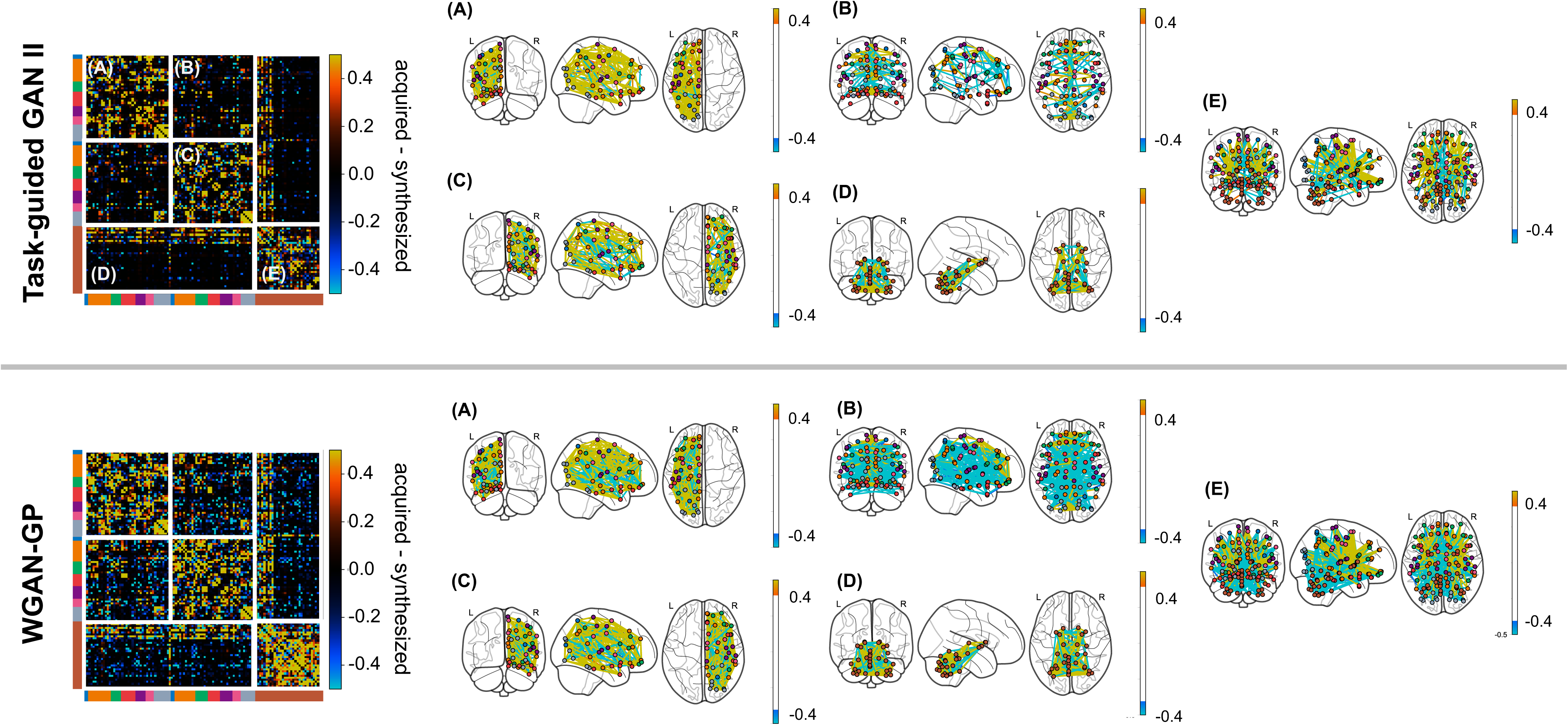
Comparison of differences between acquired and synthesized structural connectivity matrices using Task-guided GAN II and WGAN-GP. This figure illustrates the average differences across connectivity matrices, highlighting the differences between acquired and synthesized connectomes. The connectivity matrix and connectome are segmented to distinguish various connectome components: intra-hemispherical connections (left hemisphere: A, right hemisphere: C), inter-hemispherical connections (B), cerebrum-cerebellum connections (D), and intra-cerebellar connections (E).

**Figure 7** indicates a quantitative analysis of the topological features of both the acquired and synthesized connectivity matrices, utilizing graph theory metrics: connectivity strength, betweenness centrality, and clustering coefficient. The distribution of each graph metric is displayed as a density plot, along with the similarity between the distributions of the acquired and synthesized matrices estimated using KL-divergence. Notably, the KL-divergence values and density plots indicate no difference in the distribution of connectivity strength between matrices synthesized by TG GAN II and those by WGAN-GP. Conversely, the distributions of betweenness centrality and clustering coefficient suggested that TG GAN II produced matrices with topological characteristics more closely aligned to those of the acquired matrices than WGAN-GP. Smaller KL-divergence values and the similarity of density plots across all edge density thresholds evidence this.

**Figure 7.**
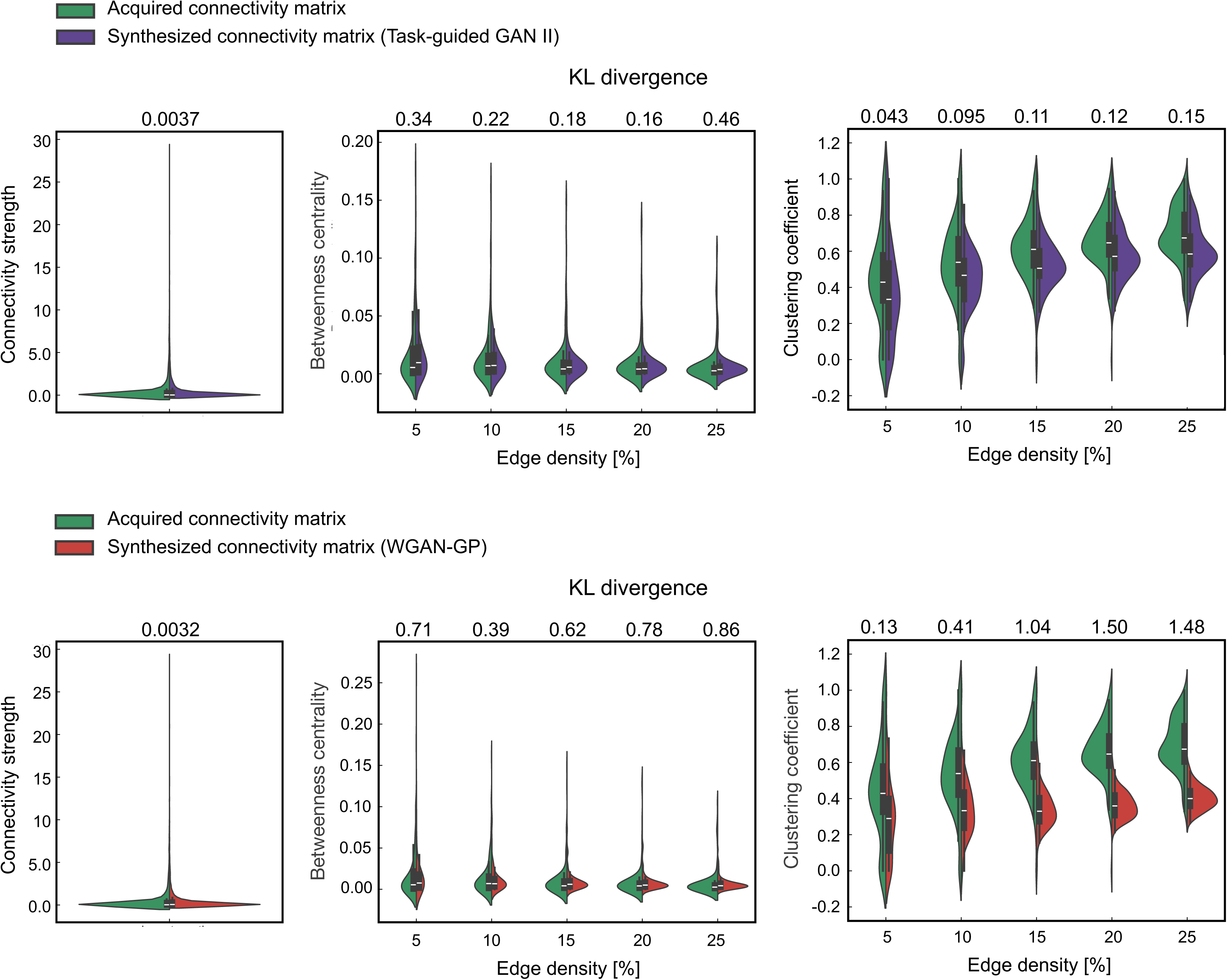
Distribution of graph theory metrics (connectivity strength, betweenness centrality, clustering coefficient) across acquired and synthesized connectivity matrices via TG GAN II and WGAN-GP under various threshold settings. The upper section displays results from Task-guided GAN II, while the lower section indicates WGAN-GP results. The similarity between the distributions of graph metrics is quantified using the Kullback-Leibler divergence.

Additionally, Figures 8 and 9 show the distribution of network properties—betweenness centrality, clustering coefficient, modularity, global efficiency, and local efficiency— calculated per-matrix basis. The effect sizes of the differences between acquired and synthesized matrices are summarized in Tables 4 and 5. Across all network measures, Task-guided GAN II exhibited lower effect sizes compared to WGAN-GP, indicating that the synthetic data generated by Task-guided GAN II more closely preserved the topological characteristics of real SC matrices.

**Figure 8.**
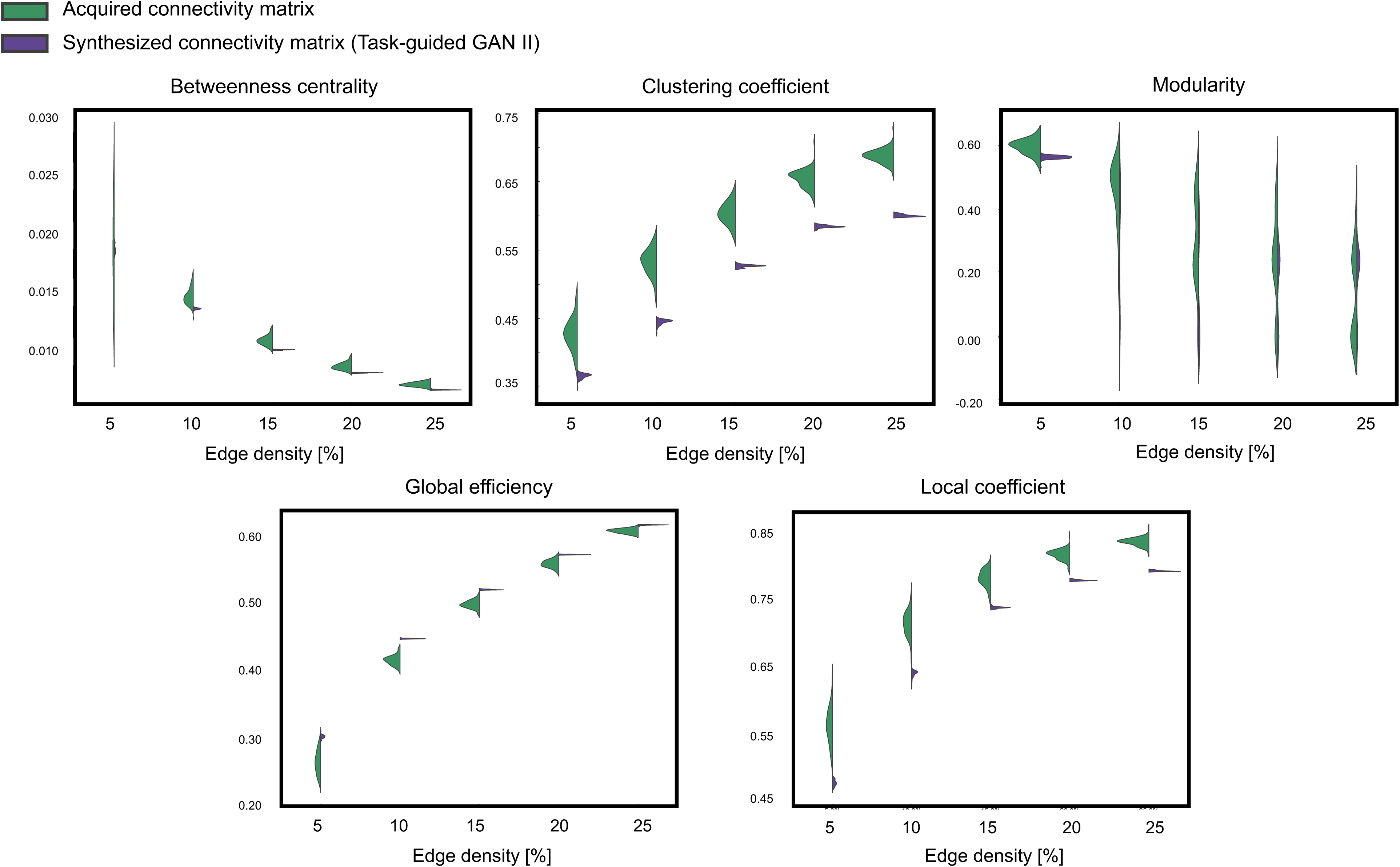
Distribution of network properties calculated on per-matrix basis across the acquired and synthesized connectivity matrices via TG GAN II.

**Figure 9.**
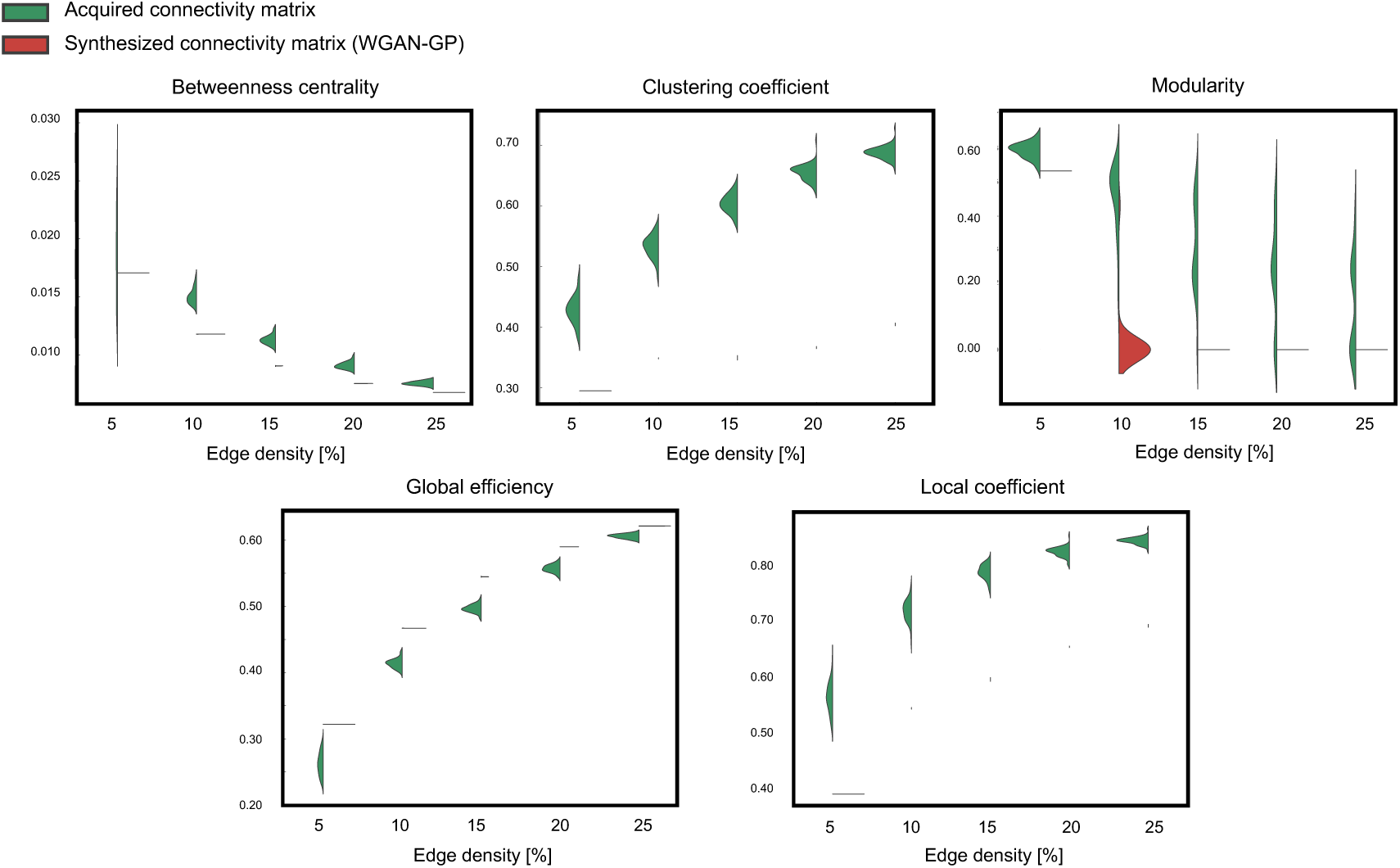
Distribution of network properties calculated on a per-matrix basis across acquired and synthesized connectivity matrices via WGAN-GP.

**Table 4.** The effect size, measured using Cohen’s d, quantifies the distributional differences in network properties between acquired and synthesized matrices. These network properties were calculated on a per-matrix basis for both real and generated data using TG GAN II.

| Edge density | Betweenness centrality | Clustering Coefficient | Modularity | Global efficiency | Local efficiency |
| --- | --- | --- | --- | --- | --- |
| 5% | -0.13 | 3.86 | 1.92 | -3.06 | 4.58 |
| 10% | 2.27 | 6.50 | 1.05 | -6.39 | 5.39 |
| 15% | 2.86 | 6.89 | 1.01 | -5.21 | 4.82 |
| 20% | 2.66 | 7.60 | 0.51 | -3.31 | 6.59 |
| 25% | 3.61 | 11.97 | -0.41 | -3.75 | 10.29 |

**Table 5.** The effect size, measured using Cohen’s d, quantifies the distributional differences in network properties between acquired and synthesized matrices. These network properties were calculated on a per-matrix basis for both real and generated data using WGAN-GP.

| Edge density | Betweenness centrality | Clustering Coefficient | Modularity | Global efficiency | Local efficiency |
| --- | --- | --- | --- | --- | --- |
| 5% | 0.59 | 8.47 | 3.74 | -4.87 | 9.24 |
| 10% | 7.04 | 14.015 | 4.49 | -10.63 | 12.79 |
| 15% | 8.13 | 23.01 | 3.05 | -11.40 | 20.82 |
| 20% | 7.48 | 31.56 | 2.11 | -7.96 | 28.85 |
| 25% | 6.40 | 39.28 | 1.36 | -6.55 | 34.97 |

### 3.2 Latent space analysis (Dataset 1)

**Figure 10** illustrates the latent spaces of each generative model, projected into two-dimensional space using PCA, with each sample annotated by its corresponding fluid intelligence score. The left side of **Figure 10** displays the PCA-projected latent spaces, while the right side shows the correlation between the first principal component (PC1) and the fluid intelligence scores. The correlation is higher in TG GAN II (*r* = 0.97, *p* < .05) than WGAN-GP (*r* = 0.15, *p* > .05) in training dataset, indicating that the latent space in TG GAN II is more closely associated with fluid intelligence scores. These results demonstrated that TG GAN II effectively captured the variance in fluid intelligence scores within its latent space. However, in the validation dataset, the correlations in both two models were non-significant (*r* = -0.25; TG GAN II, *r* = -0.23; WGAN-GP), indicating potential overfitting. These results suggest that the relationship between the latent space and fluid intelligence observed during training did not generalize to the validation dataset due to small sample size and differences in sample distributions.

**Figure 10.**
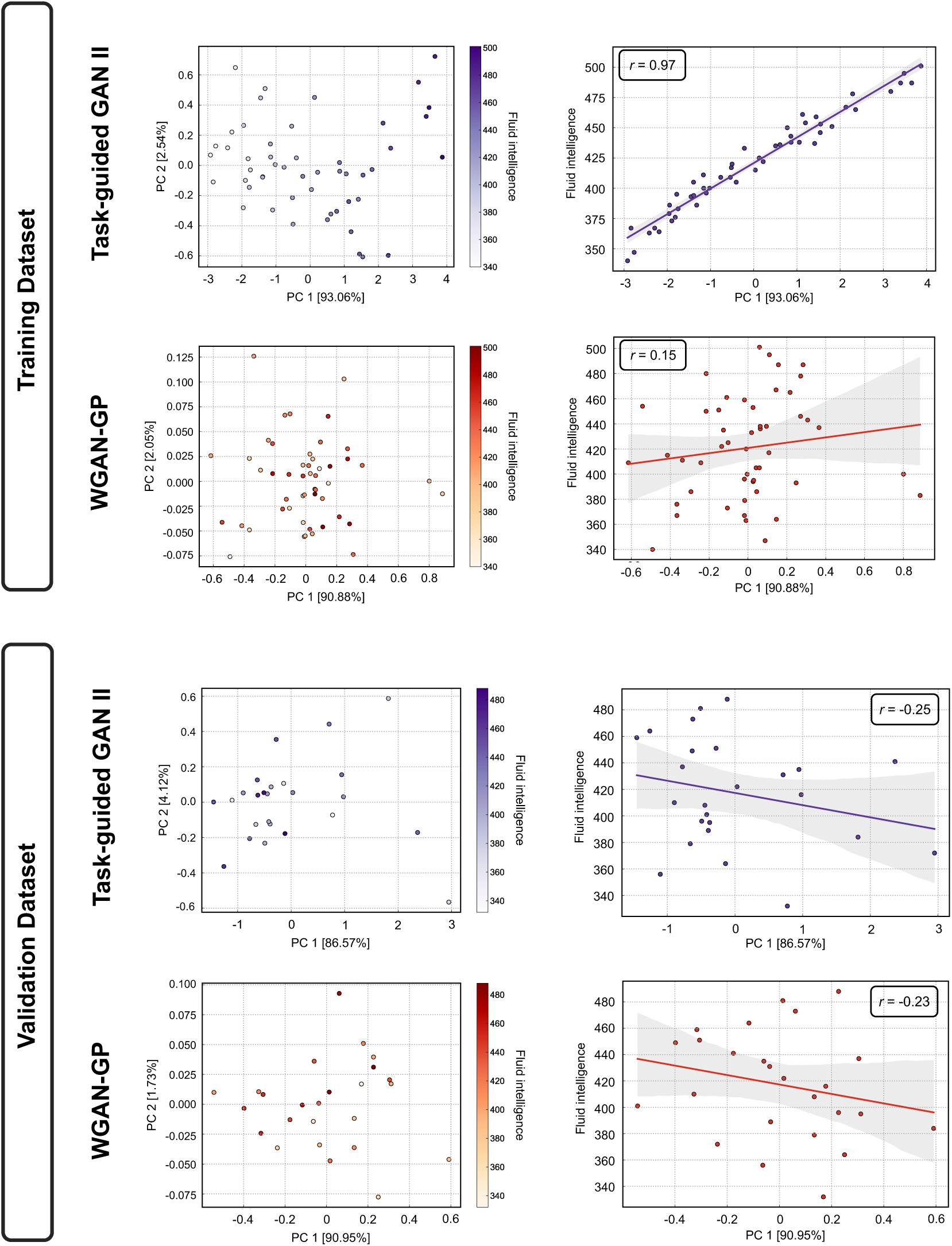
Visualization of the latent space through principal component analysis (PCA) (left) and the correlation between the first principal component (PC 1) and the fluid intelligence score (right). The latent spaces of the training dataset are shown in the upper figures, while those of the validation dataset are shown in the lower figures.

### 3.3 Prediction performance (Dataset 1)

**Figure 11** and **Figure 12** illustrate the accuracy of the prediction models using Pearson’s correlation between observed and predicted fluid intelligence scores, as well as RMSE, for data augmented by TG GAN II and WGAN-GP, respectively. The term ‘baseline’ in each figure refers to models developed solely with acquired samples without data augmentation. Each data point represents a model constructed through the repeated nested CV method. The results in **Figure 11** revealed that data augmentation with TG GAN II significantly enhanced the correlation between observed and predicted scores beyond the baseline (*p* < .01, Bonferroni-corrected), whereas augmentation with WGAN-GP did not improve. Regarding RMSE, augmentation with neither TG GAN II nor WGAN-GP led to improvements. Also, data augmentation with WGAN-GP resulted in worse RMSE than the baseline, whereas the RMSE for models augmented with TG GAN II remained consistent with the baseline (**Figure 12**). These results were consistent also in the summary of the accuracy of data-augmented and baseline prediction models indicated in **Table 6**. Notably, increasing the number of synthesized samples did not lead to further improvements in prediction accuracy, even with TG GAN II.

**Figure 11.**
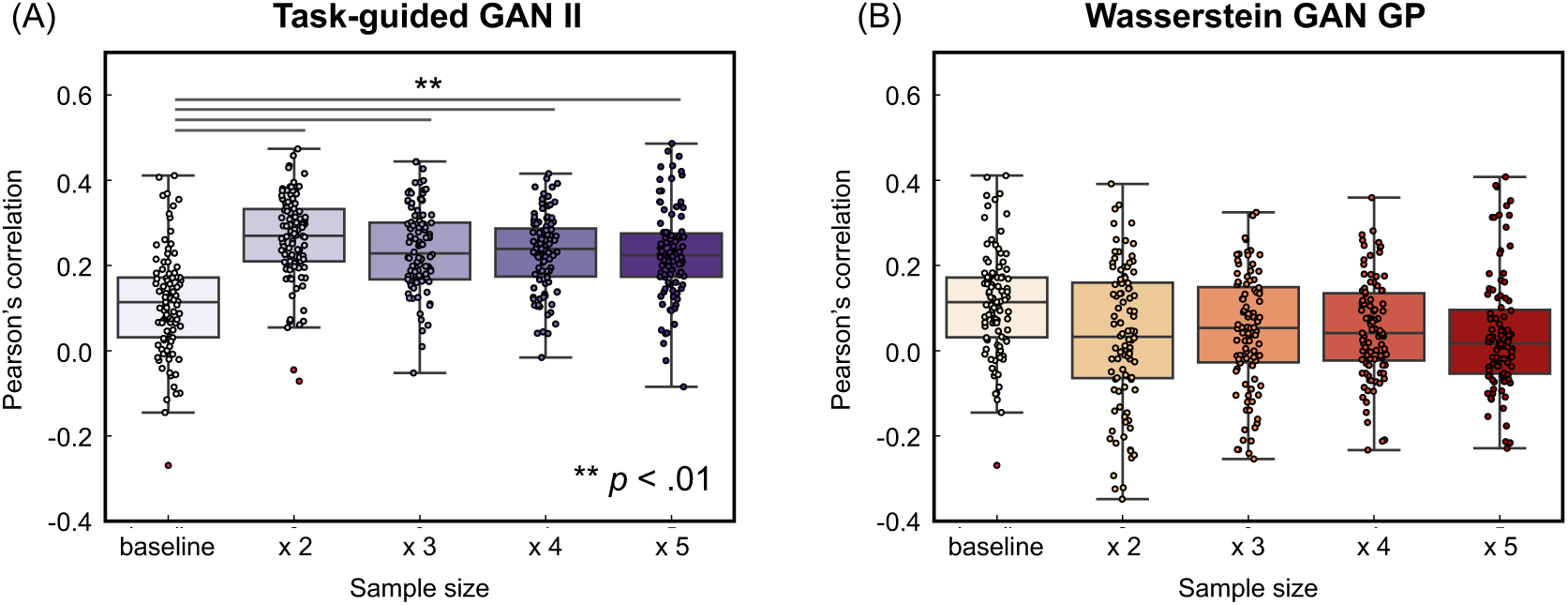
[Dataset 1] Accuracy of the prediction model, evaluated by Pearson’s correlation between observed and predicted scores. (A) indicates data augmentation results using Task-guided GAN II, while (B) presents results from employing Wasserstein GAN GP.

**Figure 12.**
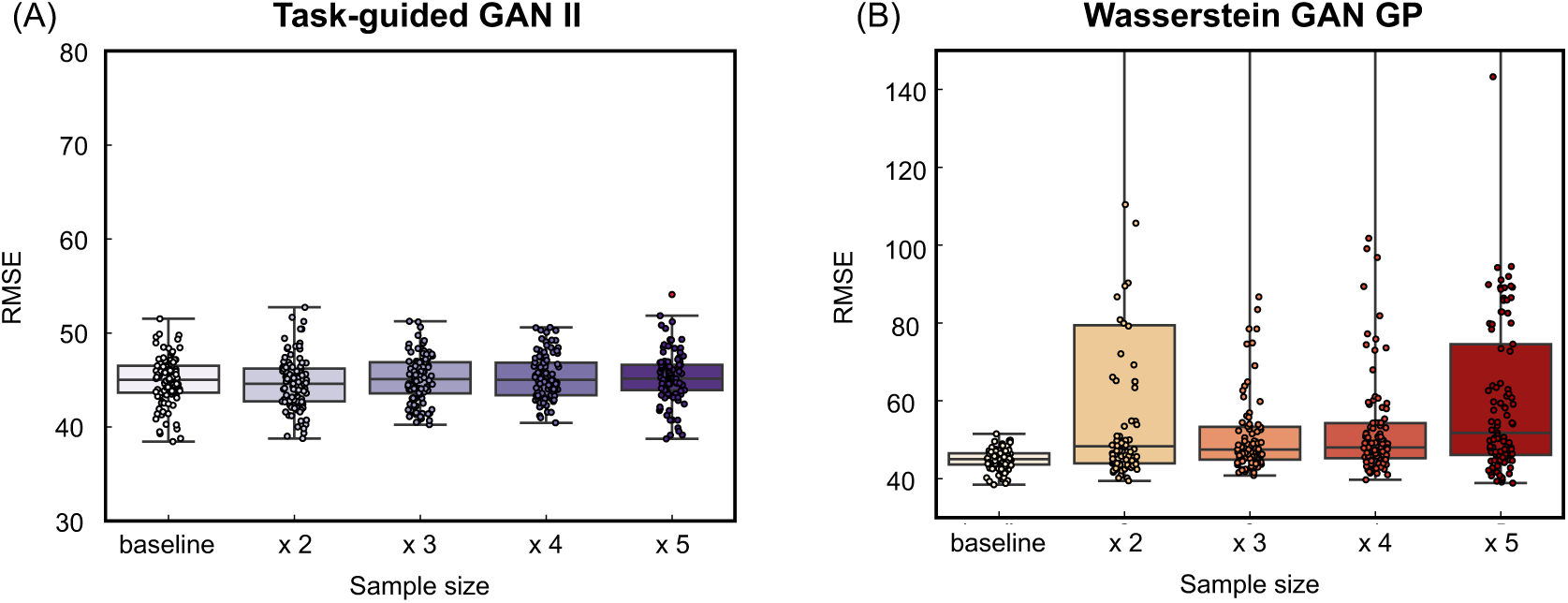
[Dataset 1] Accuracy of the prediction model, evaluated by the root mean squared error (RMSE) between observed and predicted scores. (A) indicates data augmentation results using Task-guided GAN II, while (B) presents results from employing Wasserstein GAN GP.

**Table 6.** [Dataset 1] Summary of the prediction accuracy.

| Augmentation method | Training sample size | Pearson's correlation | RMSE |
| --- | --- | --- | --- |
| Baseline | 75 | 0.11 ± 0.12 | 44.8 ± 2.53 |
| TG GAN II<br>(Proposed) | 149 (baseline + 100%) | 0.26 ± 0.10 | 44.6 ± 2.77 |
| | 223 (+ 200%) | 0.24 ± 0.10 | 45.1 ± 2.48 |
| | 297 (+ 300%) | 0.23 ± 0.09 | 45.3 ± 2.34 |
| | 371 (+ 400%) | 0.23 ± 0.11 | 45.2 ± 2.75 |
| WGAN GP | 149 (+ 100%) | 0.030 ± 0.16 | 470 ± 958 |
| | 223 (+ 200%) | 0.050 ± 0.14 | 301 ± 892 |
| | 297 (+ 300%) | 0.050 ± 0.12 | 177 ± 685 |
| | 371 (+ 400%) | 0.040 ± 0.14 | 214 ± 813 |

#### 3.3.1 Evaluation of reliability of estimated feature weight (Dataset 1)

**Figure 13** summarizes the reliability of feature weights in prediction models constructed using repeated nested cross-validation with augmented data from TG GAN II and WGAN-GP. Reliability was evaluated based on the mean and the 95% confidence interval (CI) range for each weight. A visual inspection of **Figure 13** suggests that the average feature weights are more stable across all training sample sizes when using TG GAN II for data augmentation compared to WGAN-GP. Additionally, the 95% CI for feature weights is narrower in TG GAN II than in WGAN-GP. These results indicate that prediction models augmented with TG GAN II produce more stable and reliable feature weights than those augmented with WGAN-GP.

**Figure 13.**
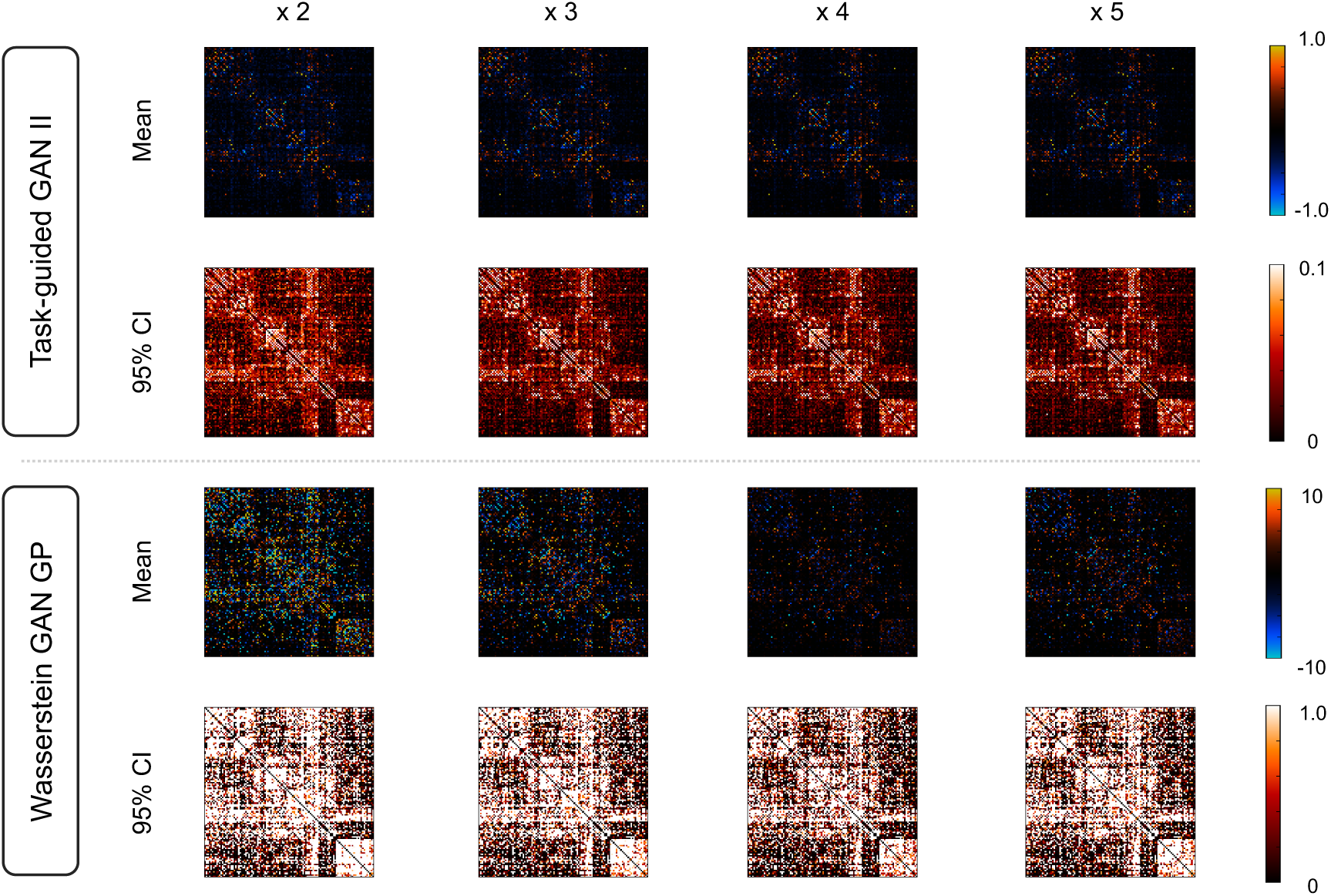
Reliability of estimated feature weights in prediction models augmented with Task-guided GAN II and Wasserstein GAN-GP, assessed based on the mean and the 95% confidence interval (CI) range for each weight.

### 3.4 Comparison of data augmentation and actual sample size increase (Dataset 2)

**Figure 14** and **Figure 15** illustrate the accuracy of the prediction models for fluid intelligence in Dataset 2, using Pearson’s correlation and RMSE between observed and predicted fluid intelligence scores. The comparison includes models trained with increasing actual sample sizes and those augmented with TG GAN II and WGAN-GP, respectively. Each data point represents a model constructed through repeated nested CV.

**Figure 14.**
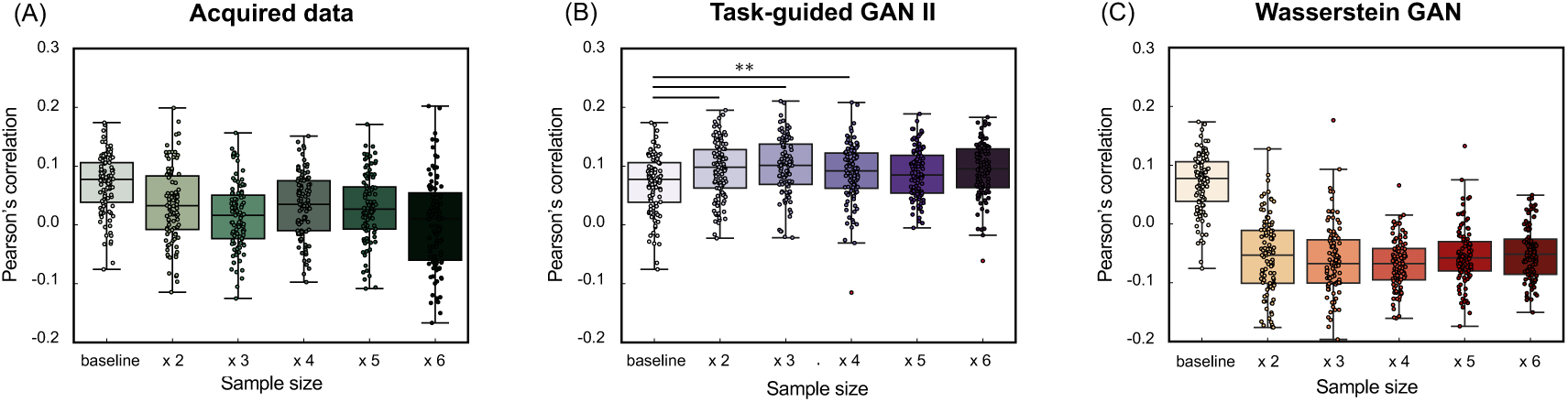
[Dataset 2] Accuracy of the prediction model for fluid intelligence, evaluated using Pearson’s correlation between observed and predicted scores. (A) shows the results of increasing the actual data sample size, (B) presents the results of data augmentation using Task-guided GAN II, and (C) displays the results from Wasserstein GAN-GP.

**Figure 15.**
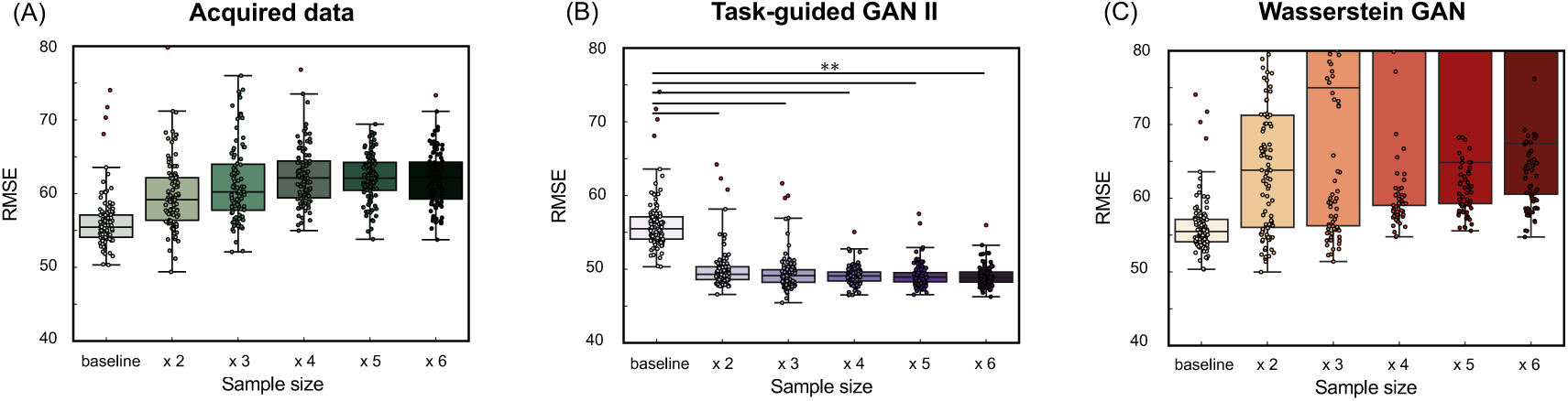
[Dataset 2] Accuracy of the prediction model for fluid intelligence, evaluated by the root mean squared error (RMSE). (A) shows the results of increasing the actual data sample size, (B) presents the results of data augmentation using Task-guided GAN II, and (C) displays the results from Wasserstein GAN-GP.

**Figure 14** shows that increasing the actual sample size improved prediction accuracy, even when using real data (A). However, data augmentation with TG GAN II significantly enhanced the correlation between observed and predicted scores compared to the baseline model (*p* < .01, Bonferroni-corrected), whereas augmentation with WGAN-GP resulted in a decline in correlation. Additionally, increasing the actual sample size and applying WGAN-GP-based data augmentation led to a deterioration in RMSE compared to the baseline (**Figure 14**). In contrast, RMSE improved in models augmented with TG GAN II. A summary of the prediction model accuracy is presented in **Table 7**.

**Table 7.** [Dataset 2] Summary of the prediction accuracy: fluid intelligence.

| Augmentation method | Training sample size | Pearson's correlation | RMSE |
| --- | --- | --- | --- |
| Baseline | 80 | $0.069 \pm 0.050$ | $56.36 \pm 3.88$ |
| Real data | 161 (baseline + 100%) | $0.034 \pm 0.063$ | $60.08 \pm 5.10$ |
| | 242 (+ 200%) | $0.015 \pm 0.056$ | $61.68 \pm 5.72$ |
| | 322 (+ 300%) | $0.033 \pm 0.054$ | $62.61 \pm 4.49$ |
| | 402 (+ 400%) | $0.029 \pm 0.057$ | $62.16 \pm 3.21$ |
| | 482 (+ 500%) | $0.0034 \pm 0.077$ | $62.17 \pm 3.72$ |
| TG GAN II<br>(Proposed) | 161 (baseline +<br>100%) | 0.094 ± 0.049 | 50.07 ± 2.77 |
| | 242 (+ 200%) | 0.099 ± 0.049 | 49.64 ± 2.59 |
| | 322 (+ 300%) | 0.088 ± 0.050 | 49.12 ± 1.29 |
| | 402 (+ 400%) | 0.087 ± 0.042 | 49.24 ± 1.62 |
| | 482 (+ 500%) | 0.093 ± 0.046 | 49.08 ± 1.42 |
| WGAN-GP | 161 (baseline +<br>100%) | -0.055 ± 0.064 | 69.07 ± 22.32 |
| | 242 (+ 200%) | -0.061 ± 0.060 | 76.18 ± 22.63 |
| | 322 (+ 300%) | -0.067 ± 0.043 | 84.64 ± 26.36 |
| | 402 (+ 400%) | -0.054 ± 0.050 | 90.38 ± 37.91 |
| | 482 (+ 500%) | -0.050 ± 0.046 | 103.32 ± 61.49 |

**Figure 16** and **Figure 17** also illustrated the accuracy of the prediction models for crystallised intelligence in Dataset 2. **Figure 16** shows that the increasing actual sample size and data augmentation with TG GAN II led to significant improvements of prediction accuracy beyond the baseline (*p* < .01, Bonferroni-corrected), whereas the augmentation with WGAN-GP did not improve. A summary of the prediction model accuracy is presented in **Table 8**.

**Figure 16.**
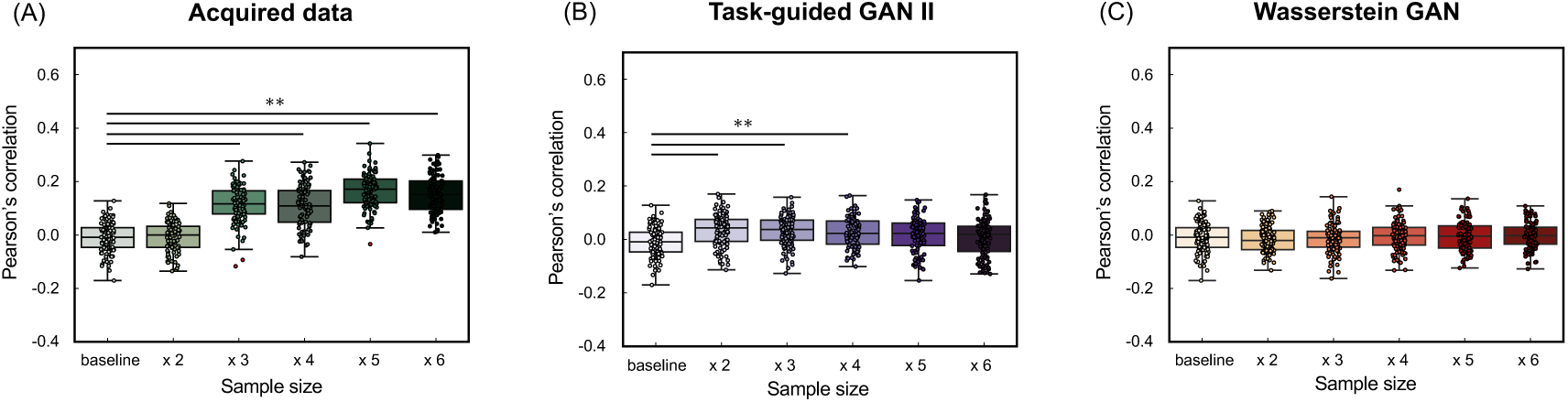
[Dataset 2] Accuracy of the prediction model for crystallised intelligence, evaluated using Pearson’s correlation between observed and predicted scores. (A) shows the results of increasing the actual data sample size, (B) presents the results of data augmentation using Task-guided GAN II, and (C) displays the results from Wasserstein GAN-GP.

**Figure 17.**
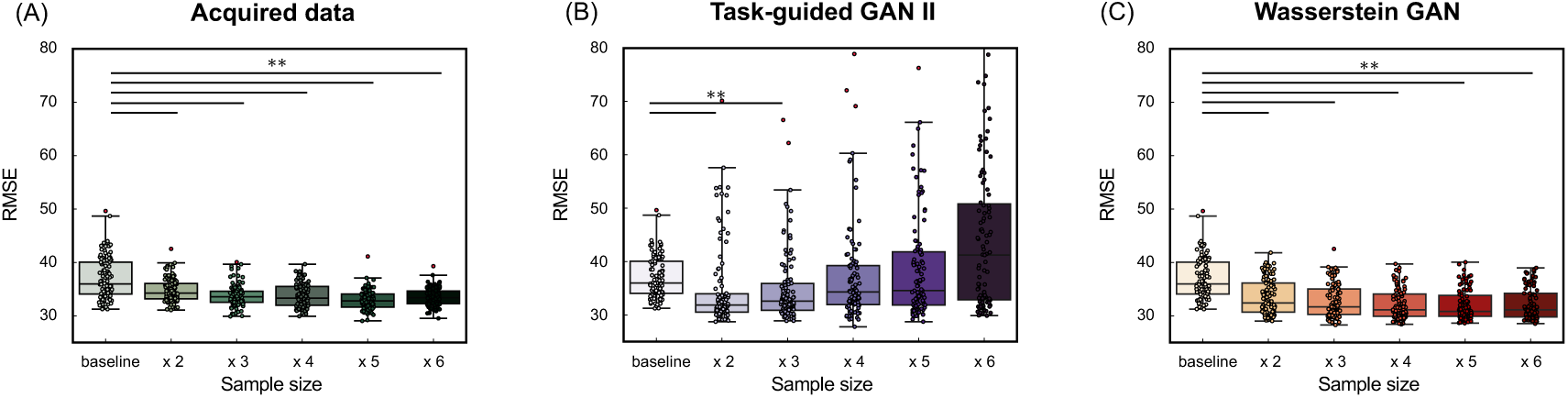
[Dataset 2] Accuracy of the prediction model for crystallised intelligence, evaluated by the root mean squared error (RMSE). (A) shows the results of increasing the actual data sample size, (B) presents the results of data augmentation using Task-guided GAN II, and (C) displays the results from Wasserstein GAN-GP.

**Table 8.** [Dataset 2] Summary of the prediction accuracy: crystallised intelligence.

| Augmentation method | Training sample size | Pearson's correlation | RMSE |
| --- | --- | --- | --- |
| Baseline | 80 | -0.010 ± 0.056 | 37.08 ± 3.89 |
| Real data | 161 (baseline + 100%) | -0.0069 ± 0.054 | 34.80 ± 2.20 |
| | 242 (+ 200%) | 0.11 ± 0.068 | 33.80 ± 2.04 |
| | 322 (+ 300%) | 0.11 ± 0.079 | 33.76 ± 2.13 |
| | 402 (+ 400%) | 0.16 ± 0.063 | 33.02 ± 1.82 |
| | 482 (+ 500%) | 0.15 ± 0.070 | 33.56 ± 1.65 |
| TG GAN II (Proposed) | 161 (baseline + 100%) | 0.034 ± 0.060 | 35.07 ± 7.74 |
| | 242 (+ 200%) | 0.031 ± 0.056 | 35.12 ± 6.78 |
| | 322 (+ 300%) | 0.026 ± 0.056 | 37.45 ± 9.15 |
| | 402 (+ 400%) | 0.015 ± 0.065 | 38.80 ± 9.64 |
| | 482 (+ 500%) | 0.0061 ± 0.069 | 44.41 ± 13.90 |
| WGAN GP | 161 (baseline + 100%) | -0.016 ± 0.059 | 33.50 ± 3.24 |
| | 242 (+ 200%) | -0.013 ± 0.054 | 32.81 ± 3.11 |
| | 322 (+ 300%) | -0.0043 ± 0.055 | 32.12 ± 2.78 |
| | 402 (+ 400%) | -0.0042 ± 0.055 | 32.04 ± 2.88 |
| | 482 (+ 500%) | -0.0022 ± 0.050 | 32.23 ± 2.95 |

## 4. Discussion

### 4.1 The relationship between graph structure and prediction accuracy

In this study, we introduced TG GAN II, a novel method for augmenting brain connectivity data, to enhance the accuracy of predictive models for human cognitive traits and behavior. The analysis of the fluid intelligence prediction model, constructed using the NIMH Healthy Research Volunteer Dataset, showed that TG GAN II could synthesize structural connectivity matrices with features more closely aligned with the actual data than its baseline model, WGAN-GP. Furthermore, data augmentation with TG GAN II resulted in improved prediction accuracy.

Upon examining the average synthesized and acquired matrices, it was observed that WGAN-GP tended to overgenerate inter-hemispherical connections compared to TG GAN II, which inaccurately reflected the fewer connections present in the actual data. Zimmermann et al. have suggested that intra-hemispherical structural connectivity is crucial for capturing the variance in cognitive traits, including fluid intelligence [37]—a notion supported by such connections in the acquired data. Unlike WGAN-GP, TG GAN II effectively minimized inter-hemispherical connections while enhancing intra-hemispherical connections, thereby contributing to the enhanced predictive accuracy of the model.

Furthermore, a graph theoretical analysis of both acquired and synthesized connectivity matrices showed that TG GAN II could produce structural connectivity matrices with topological features more closely matching those of the actual data than WGAN-GP. Garai et al. highlighted that network patterns or topological features in structural connectivity possess predictive power for human cognitive traits [38]. While Litwińczuk observed no consistent advantage in utilizing graph theory measures over connectivity values for explaining and predicting cognitive functions in healthy and typical domains, they noted instances where nodal graph theory metrics of the structural network outperformed raw connectivity models in predictive ability [39]. These prior studies suggest that the enhanced prediction accuracy observed with matrices synthesized by TG GAN II may be influenced by their graph-theoretical features, which exhibit greater similarity to those of the acquired matrices. However, further investigation is needed to determine the precise relationship between these features and prediction performance.

Consequently, our results suggest that the task-guided branch, implemented as the regression model in TG GAN II, contributed to enhancing the synthesis of relevant connectomic features for fluid intelligence prediction while suppressing ineffective features.

### 4.2 Latent space structure

The previous studies, the foundational work on Task-guided GANs [16] and BrainNetGAN [15], have demonstrated the efficacy of task-guided branches in enhancing task performance—specifically, improving age prediction accuracy from T1-weighted images and classification accuracy for Alzheimer’s disease, respectively.

In our study, by visualizing the embedded latent space using PCA, we have shown that the task-guided branch not only improved the task performance but also formed a latent space where latent variables were significantly correlated with fluid intelligence scores in the training dataset (**Figure 10**). This correlation was not observed in the validation dataset and even reversed in sign, indicating that the alignment between the latent representation and behavioral variables may not generalize across different datasets. This discrepancy suggests that, while the task-guided branch effectively promotes behavioral alignment in the latent space during training, the learned representation is prone to overfitting in generative model training. This tendency is particularly evident in scenarios with limited samples. As both the training and validation datasets were drawn from a limited-sample pool, this pattern may reflect a known limitation of representation learning under small-sample cases, where the latent space can become overfitted to sample-specific variance. Given that our method targets low-sample scenarios, such overfitting tendencies are difficult to avoid.

Furthermore, as later described in Section 4.4, the augmented samples generated from this latent space yielded improvements in prediction performance in some cases. Additionally, it is essential to note that the accuracy of the prediction models was tested on the test dataset, which is independent of the discovery dataset. This design ensures that the observed improvements in prediction performance are not merely due to overfitting in the latent representation, allowing the effectiveness of data augmentation using TG GAN II to be evaluated independently from the latent space analysis. These results suggest that even if the behavioral alignment in the latent structure does not generalize, the synthesized samples through the trained generative models may still be useful for training predictive models with limited samples. Moreover, we observed a similar pattern of correlation reversal in the validation dataset of the WGAN-GP model, which does not include the task-guided branch. This suggests that the instability of latent–behavior alignment may not be specific to our task-guided framework but rather could reflect a broader limitation of generative modeling under low-sample conditions.

Notably, prior work on task-guided generative models has not explicitly examined the correlation between latent variables and behavioral outcomes. By applying PCA to the latent space and quantifying its alignment with cognitive scores, our study provided a first attempt to link task-guided training of the generative model with interpretable latent structures. This approach offers an initial step toward understanding how outcome variables are encoded in latent space and establishes a foundation for future research on interpretable generative modeling in neuroimaging.

### 4.3 Reliability of feature weights

We investigated the reliability of feature weights in prediction models with data augmentation as the aim to predict the human cognitive traits is for identifying the high predictive and important neural connection in human cognitions [40]. In functional connectivity-based predictive modeling, Chen et al. stated that low variance in feature importance (i.e. coefficient calculated in a prediction model) is necessary for identifying truly important connectivity in a cognition [41].

In Dataset 1, we examined the stability of feature weights using the mean and 95% CI of predicted feature weights in the repeated nested CV folds. These results showed that the variation of mean across training sample sizes and 95% CI in data augmented model using TG GAN II were smaller than WGAN-GP (**Figure 13**). It indicates that estimated feature weights in prediction model with TG GAN II-based data augmentation are stable compared with WGAN-GP. The out-of-sample prediction performance was also better in TG GAN-II than WGAN-GP, as shown in **Figure 11**. These findings suggest that predictive features can be identified and sharpened in prediction models using data augmentation with TG GAN II. However, as stated in Chen et al., we should be careful that low variance of estimated features is necessary but not sufficient for validating true predictive connectivity [40].

### 4.4 Can data augmentation serve as a substitute for increasing the actual dataset size?

This study investigated whether GAN-based data augmentation can effectively substitute for increasing real data size in structural connectivity-based cognitive prediction. The results of fluid intelligence prediction indicated that increasing the real dataset size did not significantly improve the prediction accuracy. Pearson’s correlation remained close to zero, and RMSE showed minimal improvement even as the sample size increased.

In contrast, TG GAN II-based data augmentation contributed to improved prediction accuracy. The RMSE decreased as the augmentation ratio increased, with the lowest RMSE (49.08) observed at 500% augmentation, compared to 56.36 in the baseline model (**Table 7**). However, although the RMSE improved, correlation coefficients did not show a consistent trend of improvement with increasing augmentation levels. This suggests that while TG GAN II successfully captured meaningful structural connectivity patterns, its ability to introduce diversity beneficial to prediction is limited beyond a certain augmentation threshold.

Conversely, WGAN-GP-based augmentation severely degraded prediction performance. Across all augmentation levels, Pearson’s correlation remained negative, and RMSE exceeded 100, indicating that standard GANs failed to preserve connectivity structures critical for fluid intelligence prediction.

On the other hand, unlike fluid intelligence prediction, increasing the real dataset size substantially improved crystallized intelligence prediction accuracy. Pearson’s correlation increased from -0.010 (baseline) to 0.15 (482 samples), while RMSE decreased from 37.08 to 33.56, demonstrating that increasing the real dataset effectively enhances model performance for crystallized intelligence prediction (**Table 8**).

The effect of TG GAN II-based augmentation on crystallized intelligence prediction was more limited compared to fluid intelligence. Augmenting the dataset up to 300% led to modest improvements in RMSE and correlation but further increases in augmentation ratio resulted in performance deterioration, with RMSE increasing to 44.41 at 500% augmentation. This trend suggests that GAN-based augmentation may not effectively capture the underlying patterns necessary for crystallized intelligence prediction, particularly at higher augmentation levels.

Interestingly, WGAN-GP showed comparable RMSE values to real data in crystallized intelligence prediction, although it did not improve correlation. This contrasts with the results observed for fluid intelligence, where WGAN-GP severely degraded prediction performance. One possible interpretation is that the structural properties of the synthesized connectivity matrices were more similar to real data for crystallized intelligence. However, it remains unclear whether these synthesized samples adequately captured the relevant variability necessary for behavioral prediction. Further analysis is needed to determine how WGAN-GP preserves meaningful structural and behavioral relationships in different cognitive traits.

These results indicate that GAN-based data augmentation cannot fully replace increasing real data size. Although augmentation was beneficial within a restricted range (up to 200– 300%), excessive augmentation led to saturation of performance improvement and, in some cases, performance degradation. The effectiveness of augmentation depends on the information content of the baseline dataset. If important predictive features are absent in the baseline dataset but present in newly acquired real data, increasing real data would be a more effective strategy than augmentation.

Given these findings, data augmentation should be considered a complementary tool rather than a full substitute for real data collection. In scenarios where acquiring large datasets (e.g., HCP) is impractical, GAN-based augmentation may provide an alternative means to expand training datasets and improve model robustness. However, careful tuning of augmentation ratios is necessary to avoid overfitting and ensure that synthetic data contributes meaningfully to model generalization.

The results from Dataset 1 demonstrated that augmentation was effective, whereas Dataset 2 indicated that augmentation was beneficial only up to 200% expansion in our experiments. This suggests that the effectiveness of augmentation may vary depending on dataset characteristics and the regression model used. Future studies should explore how different datasets, cognitive traits, and prediction model architectures interact to determine the optimal conditions for augmentation.

Despite these limitations, the introduction of the TG GAN II in this study represents a significant contribution to network neuroscience by providing a novel tool for augmenting connectivity matrix data. By incorporating a task-guided branch into the standard GAN architecture, TG GAN II enhances the generation of synthetic structural connectivity matrices, offering a potential solution for improving predictive modeling in scenarios with limited real data. This approach expands the methodological varieties available for network neuroscience research, enabling further exploration of generative modeling techniques for brain connectivity analysis.

### 4.5 Limitations and future work

The demonstrated efficacy of our TG GAN II in connectome-based prediction tasks suggested future directions for further investigations despite some limitations that need discussion.

While our evaluation focused on synthesizing structural connectomes, the versatile framework of TG GAN II is not confined to this application alone; it holds the potential for enhancing functional connectome-based predictions through data augmentation. In future work, we will explore whether TG GAN II’s augmentation capabilities can indeed improve the accuracy of predictions based on functional connectomes.

Additionally, this study focused on intelligence as the target prediction variable. However, previous research has explored other cognitive and behavioral traits, such as motor function and creativity. Future studies will validate the effectiveness of TG GAN II in augmenting data for the prediction of these and other behavioral traits.

Moreover, while the present study assessed the impact of data augmentation on prediction accuracy, future work should evaluate the generalization performance of regression models trained with augmented data using an independent dataset. This will help determine whether synthetic samples truly enhance model robustness and transferability across different populations.

Furthermore, this study proposed a GAN-based data augmentation framework for connectivity-based prediction. However, further refinements to generative models are necessary to enhance their practicality. For example, the base generative model could be pretrained on a large, well-established dataset and subsequently fine-tuned or adapted via transfer learning for use with smaller sample-size datasets, improving its generalizability and robustness.

Finally, we attempted to understand the representational structure introduced by the task-guided branch in generative models by visualizing and analyzing the correlation between the latent variables of TG GAN II and fluid intelligence. However, the strong correlation observed in the training data was not replicated in the validation data, suggesting potential overfitting in generative model training. Nevertheless, data augmentation using the proposed model improved prediction accuracy under specific conditions, indicating that a strong correlation between latent variables and behavioral traits may not be a prerequisite for effective data augmentation. In future work, it will be essential to test other analytical methods for characterizing the latent space structure, as well as to gain a deeper understanding of how task-guided mechanisms influence the learned latent representations. These directions would be beneficial for elucidating why the task-guided branch contributes to improved prediction performance in data augmentation.

## 5. Acknowledgements

This work was supported by JSPS KAKENHI Grant Number JP24K03028.

## 7. Supporting Information

The source code for the methods used in this paper will be made available at https://github.com/MIS-Lab-Doshisha/tg-gan2.

## References

[1] O. Sporns, G. Tononi, and R. Kötter, “The Human Connectome: A Structural Description of the Human Brain,” PLOS Computational Biology, vol. 1, no. 4, p. e42, Sep. 2005, doi: 10.1371/journal.pcbi.0010042.

[2] O. Sporns, “Structure and function of complex brain networks,” Dialogues in Clinical Neuroscience, vol. 15, no. 3, pp. 247–262, Sep. 2013, doi: 10.31887/DCNS.2013.15.3/osporns.

[3] X. Shen et al., “Using connectome-based predictive modeling to predict individual behavior from brain connectivity,” Nat Protoc, vol. 12, no. 3, Art. no. 3, Mar. 2017, doi: 10.1038/nprot.2016.178.

[4] D. Scheinost et al., “Ten simple rules for predictive modeling of individual differences in neuroimaging,” NeuroImage, vol. 193, pp. 35–45, Jun. 2019, doi: 10.1016/j.neuroimage.2019.02.057.

[5] A. W. K. Yeung, S. More, J. Wu, and S. B. Eickhoff, “Reporting details of neuroimaging studies on individual traits prediction: A literature survey,” NeuroImage, vol. 256, p. 119275, Aug. 2022, doi: 10.1016/j.neuroimage.2022.119275.

[6] Z. Cui and G. Gong, “The effect of machine learning regression algorithms and sample size on individualized behavioral prediction with functional connectivity features,” NeuroImage, vol. 178, pp. 622–637, Sep. 2018, doi: 10.1016/j.neuroimage.2018.06.001.

[7] D. C. Van Essen, S. M. Smith, D. M. Barch, T. E. J. Behrens, E. Yacoub, and K. Ugurbil, “The WU-Minn Human Connectome Project: An overview,” NeuroImage, vol. 80, pp. 62–79, Oct. 2013, doi: 10.1016/j.neuroimage.2013.05.041.

[8] N. D. Volkow et al., “The conception of the ABCD study: From substance use to a broad NIH collaboration,” Developmental Cognitive Neuroscience, vol. 32, pp. 4– 7, Aug. 2018, doi: 10.1016/j.dcn.2017.10.002.

[9] C. Sudlow et al., “UK Biobank: An Open Access Resource for Identifying the Causes of a Wide Range of Complex Diseases of Middle and Old Age,” PLOS Medicine, vol. 12, no. 3, p. e1001779, Mar. 2015, doi: 10.1371/journal.pmed.1001779.

[10] A. R. Laird, “Large, open datasets for human connectomics research: Considerations for reproducible and responsible data use,” NeuroImage, vol. 244, p. 118579, Dec. 2021, doi: 10.1016/j.neuroimage.2021.118579.

[11] L. K. Avberšek and G. Repovš, “Deep learning in neuroimaging data analysis: Applications, challenges, and solutions,” Frontiers in Neuroimaging, vol. 1, 2022, Accessed: Jul. 18, 2023. [Online]. Available: https://www.frontiersin.org/articles/10.3389/fnimg.2022.981642

[12] B. Barile, A. Marzullo, C. Stamile, F. Durand-Dubief, and D. Sappey-Marinier, “Data augmentation using generative adversarial neural networks on brain structural connectivity in multiple sclerosis,” Computer Methods and Programs in Biomedicine, vol. 206, p. 106113, Jul. 2021, doi: 10.1016/j.cmpb.2021.106113.

[13] M. S. Meor Yahaya and J. Teo, “Data augmentation using generative adversarial networks for images and biomarkers in medicine and neuroscience,” Frontiers in Applied Mathematics and Statistics, vol. 9, 2023, Accessed: May 21, 2023. [Online]. Available: https://www.frontiersin.org/articles/10.3389/fams.2023.1162760

[14] Q. Zuo et al., “Hemisphere-Separated Cross-Connectome Aggregating Learning via VAE-GAN for Brain Structural Connectivity Synthesis,” IEEE Access, vol. 11, pp. 48493–48505, 2023, doi: 10.1109/ACCESS.2023.3276989.

[15] C. Li, Y. Wei, X. Chen, and C.-B. Schönlieb, “BrainNetGAN: Data Augmentation of Brain Connectivity Using Generative Adversarial Network for Dementia Classification,” in Deep Generative Models, and Data Augmentation, Labelling, and Imperfections, S. Engelhardt, I. Oksuz, D. Zhu, Y. Yuan, A. Mukhopadhyay, N. Heller, S. X. Huang, H. Nguyen, R. Sznitman, and Y. Xue, Eds., Cham: Springer International Publishing, 2021, pp. 103–111. doi: 10.1007/978-3-030-88210-5_9.

[16] R. Li, M. Bastiani, D. Auer, C. Wagner, and X. Chen, “Image Augmentation Using a Task Guided Generative Adversarial Network for Age Estimation on Brain MRI,” in Medical Image Understanding and Analysis, B. W. Papież, M. Yaqub, J. Jiao, A. I. L. Namburete, and J. A. Noble, Eds., Cham: Springer International Publishing, 2021, pp. 350–360. doi: 10.1007/978-3-030-80432-9_27.

[17] Y.-F. Tan, C.-M. Ting, F. Noman, R. C.-W. Phan, and H. Ombao, “Graph-Regularized Manifold-Aware Conditional Wasserstein GAN for Brain Functional Connectivity Generation,” Dec. 10, 2022, arXiv: arXiv:2212.05316. doi: 10.48550/arXiv.2212.05316.

[18] I. J. Goodfellow, et al., “Generative Adversarial Networks,” Jun. 10, 2014, arXiv: arXiv:1406.2661.doi: 10.48550/arXiv.1406.2661.

[19] A. C. Nugent et al., “The NIMH intramural healthy volunteer dataset: A comprehensive MEG, MRI, and behavioral resource,” Sci Data, vol. 9, no. 1, Art. no. 1, Aug. 2022, doi: 10.1038/s41597-022-01623-9.

[20] I. Gulrajani, F. Ahmed, M. Arjovsky, V. Dumoulin, and A. C. Courville, “Improved Training of Wasserstein GANs,” in Advances in Neural Information Processing Systems, Curran Associates, Inc., 2017. Accessed: Feb. 08, 2024. [Online]. Available: https://proceedings.neurips.cc/paper_files/paper/2017/hash/892c3b1c6dccd52936e 27cbd0ff683d6-Abstract.html

[21] J. Kawahara et al., “BrainNetCNN: Convolutional neural networks for brain networks; towards predicting neurodevelopment,” NeuroImage, vol. 146, pp. 1038– 1049, Feb. 2017, doi: 10.1016/j.neuroimage.2016.09.046.

[22] R. C. Gershon, M. V. Wagster, H. C. Hendrie, N. A. Fox, K. F. Cook, and C. J. Nowinski, “NIH Toolbox for Assessment of Neurological and Behavioral Function,” Neurology, vol. 80, no. 11 Supplement 3, pp. S2–S6, Mar. 2013, doi: 10.1212/WNL.0b013e3182872e5f.

[23] R. K. Heaton et al., “Reliability and Validity of Composite Scores from the NIH Toolbox Cognition Battery in Adults,” Journal of the International Neuropsychological Society, vol. 20, no. 6, pp. 588–598, Jul. 2014, doi: 10.1017/S1355617714000241.

[24] M. Cieslak et al., “QSIPrep: an integrative platform for preprocessing and reconstructing diffusion MRI data,” Nat Methods, vol. 18, no. 7, pp. 775–778, Jul. 2021, doi: 10.1038/s41592-021-01185-5.

[25] J. Veraart, D. S. Novikov, D. Christiaens, B. Ades-aron, J. Sijbers, and E. Fieremans, “Denoising of diffusion MRI using random matrix theory,” Neuroimage, vol. 142, pp. 394–406, Nov. 2016, doi: 10.1016/j.neuroimage.2016.08.016.

[26] J. L. R. Andersson, S. Skare, and J. Ashburner, “How to correct susceptibility distortions in spin-echo echo-planar images: application to diffusion tensor imaging,” Neuroimage, vol. 20, no. 2, pp. 870–888, Oct. 2003, doi: 10.1016/S1053-8119(03)00336-7.

[27] J. L. R. Andersson and S. N. Sotiropoulos, “An integrated approach to correction for off-resonance effects and subject movement in diffusion MR imaging,” Neuroimage, vol. 125, pp. 1063–1078, Jan. 2016, doi: 10.1016/j.neuroimage.2015.10.019.

[28] T. Dhollander, R. Mito, and A. Connelly, “Single-Shell 3-Tissue CSD (SS3T-CSD) modelling of developing HCP (dHCP) diffusion MRI data,” Jun. 2019.

[29] T. Dhollander, D. Raffelt, and A. Connelly, “Unsupervised 3-tissue response function estimation from single-shell or multi-shell diffusion MR data without a co-registered T1 image,” Sep. 2016.

[30] J.-D. Tournier, F. Calamante, and A. Connelly, “Improved probabilistic streamlines tractography by 2nd order integration over fibre orientation distributions”.

[31] R. E. Smith, J.-D. Tournier, F. Calamante, and A. Connelly, “Anatomically-constrained tractography: Improved diffusion MRI streamlines tractography through effective use of anatomical information,” NeuroImage, vol. 62, no. 3, pp. 1924–1938, Sep. 2012, doi: 10.1016/j.neuroimage.2012.06.005.

[32] R. E. Smith, J.-D. Tournier, F. Calamante, and A. Connelly, “SIFT2: Enabling dense quantitative assessment of brain white matter connectivity using streamlines tractography,” Neuroimage, vol. 119, pp. 338–351, Oct. 2015, doi: 10.1016/j.neuroimage.2015.06.092.

[33] Z. Cui, M. Su, L. Li, H. Shu, and G. Gong, “Individualized Prediction of Reading Comprehension Ability Using Gray Matter Volume,” Cerebral Cortex, vol. 28, no. 5, pp. 1656–1672, May 2018, doi: 10.1093/cercor/bhx061.

[34] A. E. Hoerl and R. W. Kennard, “Ridge Regression: Biased Estimation for Nonorthogonal Problems,” Technometrics, vol. 12, no. 1, pp. 55–67, Feb. 1970, doi: 10.1080/00401706.1970.10488634.

[35] R. M. Birn et al., “The effect of scan length on the reliability of resting-state fMRI connectivity estimates,” NeuroImage, vol. 83, pp. 550–558, Dec. 2013, doi: 10.1016/j.neuroimage.2013.05.099.

[36] Z. Cui, M. Su, L. Li, H. Shu, and G. Gong, “Individualized Prediction of Reading Comprehension Ability Using Gray Matter Volume,” Cereb Cortex, vol. 28, no. 5, pp. 1656–1672, May 2018, doi: 10.1093/cercor/bhx061.

[37] J. Zimmermann, J. D. Griffiths, and A. R. McIntosh, “Unique Mapping of Structural and Functional Connectivity on Cognition,” J. Neurosci., vol. 38, no. 45, pp. 9658– 9667, Nov. 2018, doi: 10.1523/JNEUROSCI.0900-18.2018.

[38] S. Garai, F. Xu, D. A. Duong-Tran, Y. Zhao, and L. Shen, “Mining Correlation between Fluid Intelligence and Whole-brain Large Scale Structural Connectivity,” AMIA Jt Summits Transl Sci Proc, vol. 2023, pp. 225–233, 2023.

[39] M. C. Litwińczuk, N. Muhlert, N. Trujillo-Barreto, and A. Woollams, “Using graph theory as a common language to combine neural structure and function in models of healthy cognitive performance,” Hum Brain Mapp, vol. 44, no. 8, pp. 3007–3022, Jun. 2023, doi: 10.1002/hbm.26258.

[40] J. Chen et al., “Relationship between prediction accuracy and feature importance reliability: An empirical and theoretical study,” NeuroImage, vol. 274, p. 120115, Jul. 2023, doi: 10.1016/j.neuroimage.2023.120115.

[41] Y. Tian and A. Zalesky, “Machine learning prediction of cognition from functional connectivity: Are feature weights reliable?,” NeuroImage, vol. 245, p. 118648, Dec. 2021, doi: 10.1016/j.neuroimage.2021.118648.

